# Predictive learning induces Bayesian cognitive maps in the hippocampus

**DOI:** 10.64898/2026.06.03.729991

**Authors:** Yeowon Kim, Yul H. R. Kang

## Abstract

Navigation requires perception: location must be inferred from noisy and ambiguous egocentric sensory inputs, as in visual estimation of distance. However, many classical models of spatial representation implicitly assume that allocentric location is directly observable, thereby neglecting perceptual uncertainty. Here, we compare such a model with a Bayesian ideal observer that explicitly incorporates perceptual inference. We find that the Bayesian observer’s beliefs over location more accurately reproduce key properties of place cell activity, including place field width, area, and density, within and across environments. Using analytic arguments and numerical simulations, we show that recurrent neural networks trained to predict the next egocentric sensory input learn representations resembling Bayesian beliefs and yield place cell-like activity in both familiar and unfamiliar environments, outperforming autoencoders trained to reproduce the current input. Together, these results suggest that hippocampal circuits may construct Bayesian cognitive maps from experience through predictive perceptual learning.

## Introduction

Spatial navigation requires estimating one’s location and orientation from noisy and ambiguous sensory inputs, such as visual estimation of distance and angle relative to a wall^1^. A central concept in understanding this ability is the cognitive map: an internal representation of the environment’s structure that allows animals to localize themselves and plan trajectories even when sensory inputs are ambiguous^2,3^. Hippocampal place cells are widely believed to instantiate such a map through their spatially selective firing, with each cell exhibiting activity confined to specific allocentric regions of an environment (“place fields”)^4,5^. This internal map supports flexible navigation and adaptive behavior in complex and changing environments^6,7^.

Localization within this map relies on egocentric sensory inputs, including self-motion and visual inputs. However, both are noisy and ambiguous, introducing two distinct sources of uncertainty: transition uncertainty (from noisy movement) and perceptual uncertainty (from ambiguous sensory input). Optimal localization therefore requires integrating information across modalities and over time while accounting for uncertainty associated with each source, i.e., Bayesian inference^1,8^.

Many existing models of place fields, such as the successor representation (SR), focus on transition dynamics by modeling expected future occupancy under a movement policy^9,10^. While these models incorporate transition uncertainty, they assume that the mapping from sensory input to position is reliable, effectively ignoring perceptual uncertainty. As a result, they may fail to explain neural responses in conditions where sensory information is ambiguous or the exact shape of the environment is unknown.

In contrast, a Bayesian ideal observer explicitly accounts for both transition and perceptual uncertainty. It maintains a probability distribution over its possible locations (the posterior belief), which is updated by integrating noisy egocentric inputs with predictions from the previous time step. This model thus provides a normative framework for how the brain could construct a cognitive map under uncertainty^11,12^.

Here, we test whether place cell activity more closely reflects the SR or the beliefs of a Bayesian ideal observer. We find that the ideal observer model more accurately reproduces key features of place cell activity, especially under visual ambiguity and environmental deformations.

The next key question is how such Bayesian representations could be learned through experience. While previous studies have shown that predictive learning can give rise to allocentric spatial representations^13–18^, it remains unclear whether these representations reflect only point estimates, as demonstrated so far, or full posterior beliefs that explicitly represent perceptual uncertainty.

To address this, we trained a recurrent neural network with a predictive objective—a form of self-supervised learning—which we refer to as PreceptNet. After training, the network could predict future sensory input from past visual and self-motion inputs. Importantly, it developed internal representations that captured not only the estimated location but also uncertainty under perceptual ambiguity, even in novel environments. These representations gave rise to place cell-like activity patterns and reproduced key spatial trends observed in experimental data. As a strong control, a network trained with an autoencoding objective—another form of self-supervised learning—showed a significantly poorer match to experimental observations.

Taken together, our findings suggest that hippocampal place fields represent posterior beliefs suitable for robust localization under perceptual uncertainty, and that such representations can be learned through predictive perceptual learning.

## Results

### Models with and without perceptual uncertainty

To test the influence of perceptual uncertainty on neural representation of space, we build on established findings that perceptual inputs strongly influence the shape and distribution of place fields when the geometry of an environment is altered from a familiar one^19,20^. Although such manipulation has been shown to induce uncertainty about location in human behavioral experiments^11,21–23^, there has been no corresponding normative modeling account of how this uncertainty shapes place field structure. In addition, recent experiments by Tanni et al. show that visual input dynamically modulates place cell activity across time and space^24^, motivating us to test which computational model can best explain how sensory information and its associated uncertainty shape spatial representations. To this end, we compare three models: the SR model and two variants of the Bayesian ideal observer model.

The SR is a framework in which each state (here, an animal or agent’s location) is represented by a predictive map of future state occupancies. Rather than encoding the current state, the SR captures the expected frequency of visits to states in the future under a given policy^9,10^. In our implementation, an SR place cell tuned to a location ***ℓ*** responds in proportion to the discounted expected number of visits to ***ℓ*** starting from the animal’s current location and following the current policy (see Methods, Fig. S3). Although the SR does not represent uncertainty about the current state, SR place cells implicitly reflect transition uncertainty, as it influences the expected number of visits. However, the SR lacks a mechanism for incorporating perceptual uncertainty (Fig. 1a,b, yellow region, and Fig. 1c, top).

**Figure 1.**
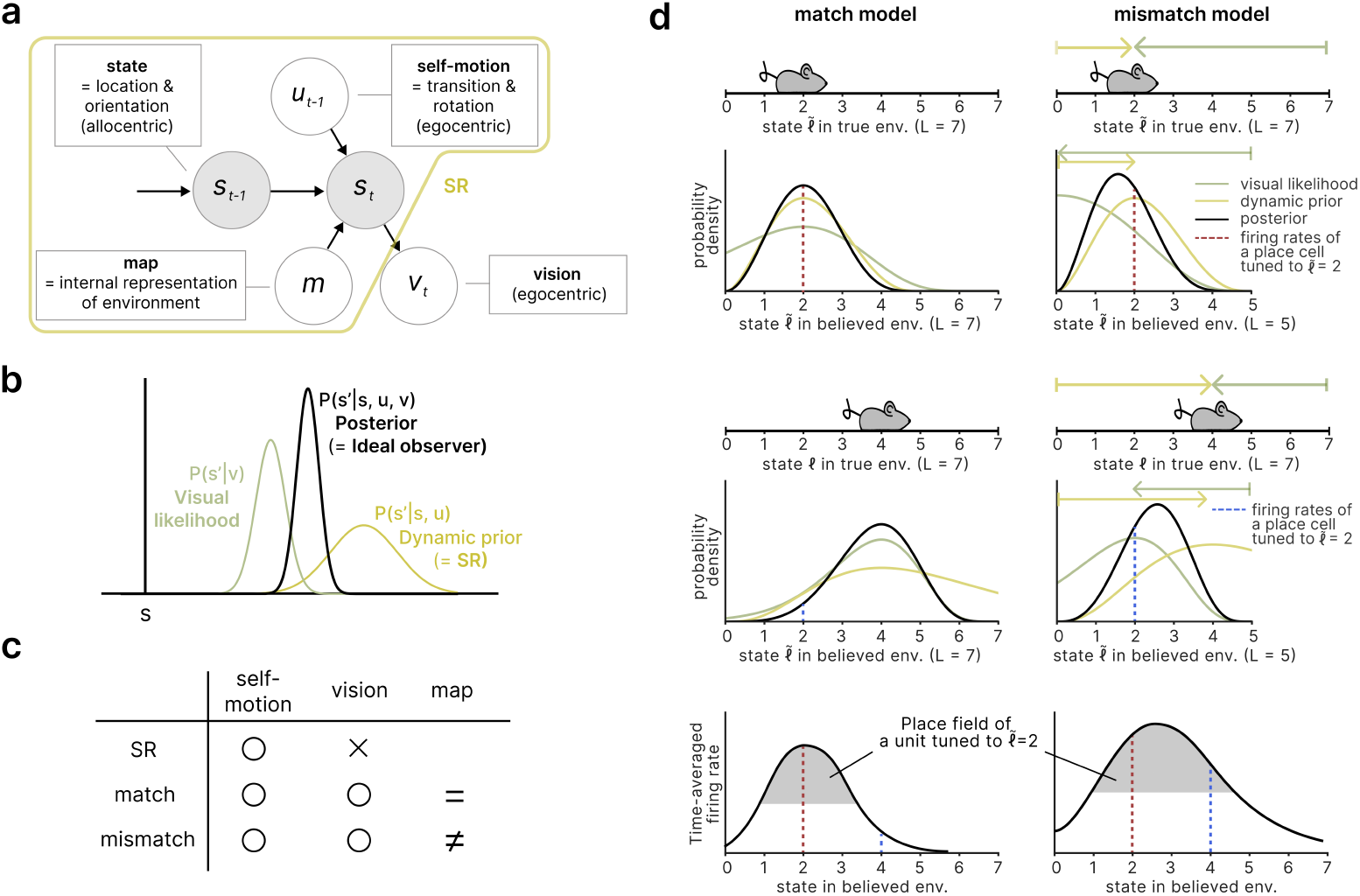
Models compared. **a**. Generative model. The animal’s state (**s**; allocentric location and orientation) is updated with the egocentric self-motion signal (**u**; translation and rotation) and determines the egocentric visual input (**v**). The state estimate also incorporates information from an internal map (**m**) representing the believed structure of the environment. The one-step SR corresponds to the same update without **v**, i.e., the prediction based solely on self-motion. **b**. Schematic of the Bayesian update. The visual likelihood (green) combines with the dynamic prior (yellow) to produce the posterior (black). **c**. Models compared. SR uses only self-motion input. The “match” model combines self-motion and visual inputs, using an internal model of the world (its “believed environment”) that matches the true environment in which the agent navigates. The “mismatch” model is identical to the match model, except that it assumes a common believed environment with size equal to the average across all true environments in the experiment, thereby mismatching each individual environment. **d**. Schematic 1D example contrasting match and mismatch model predictions (left and right, respectively). *Rows* 1 and 3. The agent’s true location (*ℓ* = 2 and *ℓ* = 4, respectively) at one time step. *Row* 2. The agent’s visual likelihood, dynamic prior, and posterior in the believed coordinates under each model. Dashed red and blue lines denote the posterior belief, which we propose to be proportional to the firing rate of a place cell tuned to 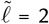 when the true location is *ℓ* = 2 and *ℓ* = 4, respectively (Rows 2, 4, and 5). For the match model, the believed environment matches the true environment (both *L* = 7), so the prior and likelihood peak near the agent’s true location, yielding a posterior that peaks at that location. For the mismatch model (believed *L* = 5), the likelihood and prior anchor to opposite walls, and the posterior lies between them. *Row* 5. Place field for the unit tuned to 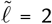, obtained by reading out the posterior probability of 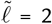 across true locations. The mismatch model’s place field is shifted and broadened depending on the tuned location and the size of the true environment (see also Fig. S2).

In contrast, the Bayesian ideal observer model infers a posterior distribution over states based on a history of noisy egocentric visual inputs and self-motion signals, employing Bayesian filtering (Fig. 1a,b). Here, “state” refers to pose, i.e., location and heading direction, whereas place fields from all models are still derived using only the location. The inference proceeds recursively in two steps: in the prediction step, the model uses self-motion signals and the previous posterior to generate a new belief about the current state, referred to as the dynamic prior. In the measurement step, the dynamic prior is refined by incorporating sensory observations through the visual likelihood, yielding the posterior over the current state. The visual likelihood is modeled as a Poisson likelihood over the full retinal image projected from the 3D environment (see Methods and Fig. 4b)^11^.

Experimental findings suggest that both components of the inference process (the dynamic prior and visual likelihood) may be represented by place cells. The role of the dynamic prior is compatible with studies showing that place cells maintain spatial tuning even in darkness, relying on self-motion and boundary cues^25–27^, while the influence of visual likelihood is compatible with the strong modulation of place cell population activity with visual inputs^28–31^. However, the posterior belief over location, which integrates the dynamic prior and visual likelihood, has not been directly compared with place fields.

Notably, the ideal observer model can operate under two distinct settings. In the *match* model, the believed environment (the agent’s internal model) matches the physical environment (Fig. 1c, middle, and Fig. 1d, left). As a result, its interpretation of sensory input reflects how the inputs are actually generated, and its beliefs remain consistent with the true distribution of its location.

However, when animals experience multiple environments, place cell activity can be anchored to walls in a particular environment^19^ or retain patterns shown in an environment across environments of partially different shapes^32–34^. This suggests that their internal model of the environment is not fully adapted to different physical environments. Therefore, we implemented this hypothesis as the *mismatch* model (Fig. 1c, bottom, and Fig. 1d, right), where the sensory input is interpreted as though it arises from a single fixed believed environment whose size equals the average across environments in the experiment. This formulation serves as a simplified approximation of partial adaptation of the internal model, rather than a claim that the internal map is strictly fixed. Under this approximation, the mismatch model serves as a principled alternative to the match model for quantitative comparison.

To better illustrate qualitative differences between the models, we examine an example neuron of an agent traveling eastward in a simple 1D environment, departing from the west wall (Fig. 1d). The mismatch model (Column 2) assumes that the agent’s believed environment is narrower than the true one. Beliefs about pose (location and heading direction) are defined within this believed environment (Rows 2 & 4). Accordingly, for a given true pose of the agent, the visual likelihood over believed locations (green curve) peaks where the believed distance to the currently faced wall equals the true distance to that wall—that is, where the observed view best matches the one predicted by the internal map (green arrows). Likewise, the dynamic prior (yellow curve) peaks where the believed distance to the wall just departed matches the agent’s true distance to that wall—that is, where the self-motion–based prediction aligns with the internal map (yellow arrows). The posterior thus lies between these wall-anchored peaks (black curve).

We posit that each place cell responds in proportion to the posterior probability of being at a location in the believed environment (its “tuned location”). Thus, the posterior over believed states is reflected in a population activity profile across neurons. To obtain a place field, we average each neuron’s activity across time steps when the agent occupied different locations in the physical environment (Fig. 1d, bottom).

Under the mismatch model, the example place field broadens and shifts toward the center of the environment because a range of physical locations, spanning front-wall-matched to rear-wall-matched positions—produce partial consistency with the same believed location (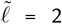; Row 5), thereby sustaining a relatively strong posterior probability at that location. In contrast, since the match model (Column 1) assumes that the believed and true environments coincide, the prior and likelihood align at the same location, producing narrow place fields centered at the true physical location.

To isolate the source of the difference between the SR and ideal observer models, we also consider a one-step variant of the SR model (Supplementary Information)^18^. In this variant, the prediction is restricted to a single step. This one-step SR corresponds to the posterior after the prediction step in the ideal observer model, that is, a model with transition uncertainty but without perceptual uncertainty: it is as though we replaced the visual input with a unique cue for each pose (One-step SR model, Fig. 1a–c). Strictly speaking, the one-step SR equals the posterior after the prediction step *plus* a delta distribution at the current location. In our implementation, we omit this delta term because it does not vary within or across environments and therefore does not influence the comparisons reported here. We verified that place fields derived from the one-step SR exhibit trends similar to those of the full SR model, indicating that the divergence between the SR and the ideal observer models arises from the inclusion of perceptual uncertainty in the latter (not shown).

### Perceptual change explains population activity change

To evaluate the models, we simulated an agent exploring four environments of increasing size (A–D; Fig. 2a) and assessed how well each model replicated key properties of place cell activity observed experimentally^24^. For the mismatch model, we fixed the believed environment across all four true environments to be an intermediate-sized environment, with width and height set to the average dimensions of the four environments.

**Figure 2.**
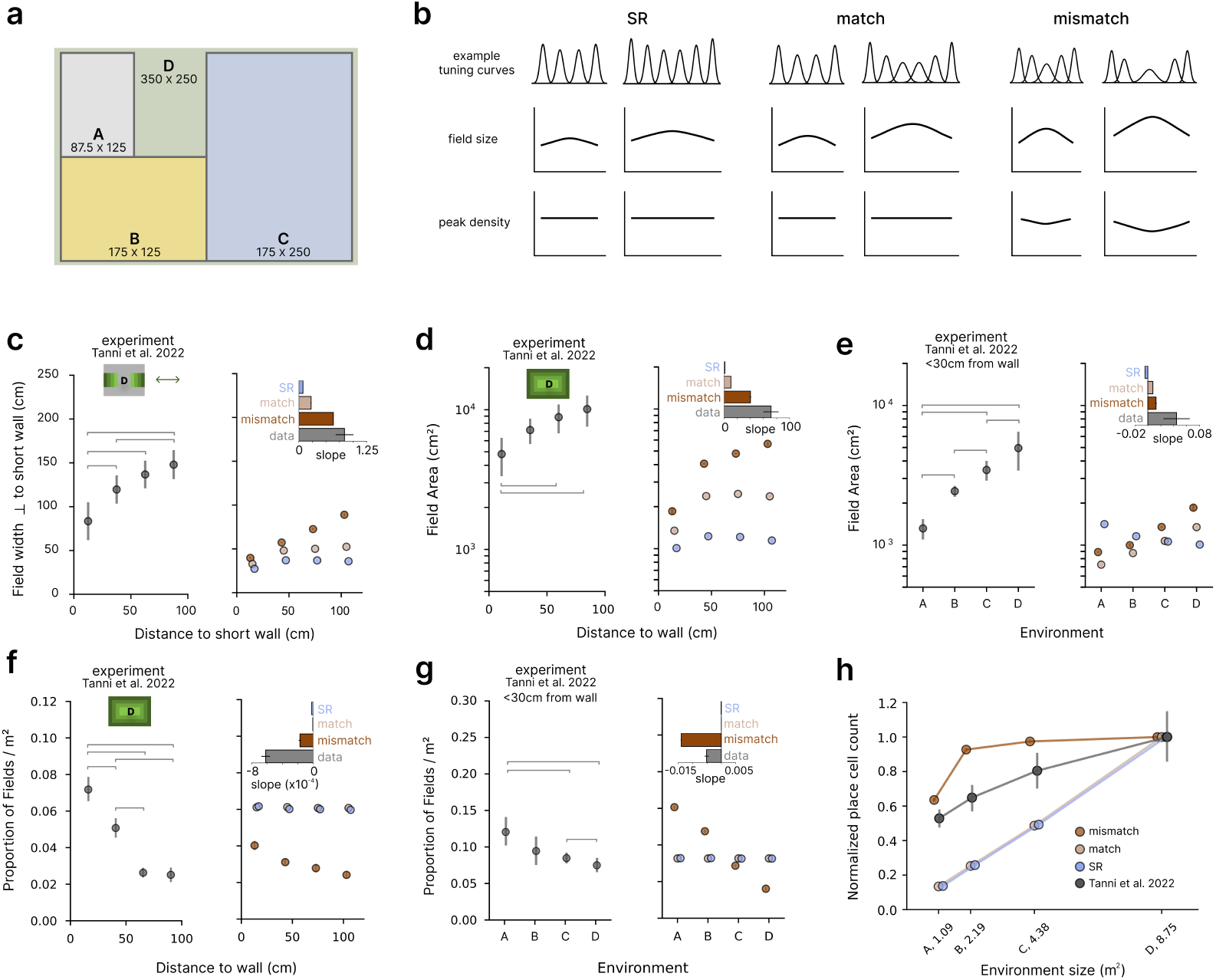
**a**. Four rectangular environments (A–D) used by Tanni et al.^24^. Numbers denote the dimensions in cm. **b**. Schematic overview of predicted place field size and density across models. Columns are organized into pairs corresponding to the SR, match, and mismatch models, with each pair showing small and large physical environments, respectively. See also Figs. S2 and S3. *Top*. Time-averaged activity profile along a 1D track. *Middle*. Place field width plotted against peak location. *Bottom*. Place field peak density plotted against location. **c-g**. Place field size and density of the data (left) and models (right). Insets show the slopes from linear regression for the data and the model predictions. Error bars show s.e.m. Horizontal bars indicate statistically significant pairwise differences in the experimental data. None of the models reproduced the degree of variability observed in the experiments, despite using different trajectories for different simulated animals (see Methods and Discussion). For this reason, we do not show significance bars in the model plots. **c**. Place field width orthogonal to the short wall as a function of distance to that wall in environment D. **d**. Place field area as a function of distance to the nearest wall in environment D. **e**. Place field area as a function of environment size. **f**. Proportion of place field peaks per unit area as a function of distance to the wall in environment D. **g**. Proportion of place field peaks per unit area as a function of environment size. **h**. Normalized number of active place cells as a function of environment size. Counts are scaled relative to the maximum value in environment D.

As described above, a core distinction between the SR and ideal observer models is that the latter explicitly incorporates perceptual uncertainty. To assess whether this distinction is reflected in population activity, we examined the correlation between moment-by-moment changes in visual input and changes in the population activity vector (Fig. S5a). Both the mismatch and match models showed robust positive correlations (mean ± SEM, *r* = 0.79 ± 0.01 and *r* = 0.79 ± 0.013), in line with experimental findings (*r* = 0.60; Fig. 5D of Ref. 24). In contrast, the SR model, which lacks access to sensory observations, showed near-zero correlation (mean ± SEM, *r* = 0.02 ± 0.01).

While this correlation highlights the role of perceptual input, it is based on the magnitude of change in the population activity vector, which relates only indirectly to spatial tuning. For instance, visual changes of opposite sign will produce the same correlation with the place cell population activity while inducing different ideal observer beliefs. To rule out this possibility and to better understand how perceptual uncertainty shapes spatial coding, we next examined how individual place fields vary across models and conditions.

### Posterior uncertainty explains place field size

To assess how place field properties vary with spatial location, we segmented each environment into nested bands based on distance to the nearest boundary (each 30 cm wide), using only the band closest to the wall when comparing across environments, as in the experimental study^24^. In both ideal observer models (match and mismatch), the posterior over location is shaped by a combination of the dynamic prior and the visual likelihood.

*Within* an environment, the posterior distribution broadens with distance from boundaries in an axis-specific manner. Uncertainty increases primarily along the axis orthogonal to the nearest wall, rather than uniformly in all directions. This directional broadening arises from the geometry of visual input—as the agent moves farther from a wall, visual cues along that axis become less informative: based on principles of pinhole optics, the Fisher information about distance *d* from the wall scales as 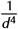 (Eq. 1 of Ref. 11). This implies that the reliability of visual input decreases sharply with distance, leading to greater uncertainty along that axis. Additionally, the posterior is broadened overall due to accumulated self-motion uncertainty in the dynamic prior, which increases with time and distance traveled.

These properties of the posterior distribution are reflected in the resulting place field structure, as schematized in Fig. 2b. Unless noted otherwise, we report the results of Kruskal-Wallis tests (KW) along with post hoc Mann-Whitney U tests with Holm-Bonferroni correction (MW) for all pairwise comparisons. Within environment D, field width increased with distance to the nearest wall along the corresponding axis in both models, but the increase was more pronounced in the mismatch model (Fig. 2c; match, KW: *H* = 16.90, *p* = 0.0007, MW: all *p <* 0.05 except 60–90 vs. 90–120 cm, n.s.; mismatch, KW: H = 17.91, *p* = 0.0005, MW: all *p <* 0.05. See also Fig. S5). This amplification in the mismatch model arises from the use of a believed environment different from the true environment, which allows multiple true (physical) locations to map onto the same believed location (Fig. 1d and Fig. S2).

In contrast, SR field width showed only a limited dependence on distance to the wall, because the transition matrix was spatially homogeneous except for truncation at the environmental boundary, resulting in relatively consistent expected future state occupancy patterns across locations (Fig. 2c; KW: *H* = 13.83, *p* = 0.003, MW: 0–30 vs. 30–60, 60–90, and 90–120 cm, *p <* 0.05; others n.s.).

These width changes translate into corresponding changes in place field area, which reflects uncertainty along both spatial axes. Accordingly, in both mismatch and match models, we observed that the average area of place fields increases with distance from the wall *within* an environment. Specifically, within environment D, field size increased with distance from the wall (Fig. 2d, right; match, KW: *H* = 15.83, *p* = 0.001, MW: all *p <* 0.05 except 30–60 vs. 90–120 cm, n.s.; mismatch, KW: *H* = 17.86, *p* = 0.0005, MW: all *p <* 0.05; see also Fig. S4c,d for environments B and C). Additionally, across environments, as larger environments include more locations far from boundaries, place field size increased with environment size in both models (Fig. 2e, right; mismatch, KW: *H* = 17.86, *p* = 0.0005, MW: all *p <* 0.05; match, KW: *H* = 17.86, *p* = 0.0005, MW: all *p <* 0.05).

In contrast, *within* each environment, SR field area showed a weaker dependence on distance to the wall, consistent with the transition matrix being approximately spatially homogeneous (area: Fig. 2d; KW: *H* = 16.21, *p* = 0.001, MW: all *p <* 0.05 except 30–60 vs. 60–90 cm, n.s.). *Across* environments, the SR model predicts the opposite trend: place field size decreases with increasing environment size (Fig. 2e; KW: *H* = 17.58, *p* = 0.0005, MW: all *p <* 0.05), because smaller environments constrain movement, increasing the frequency of state revisits and broadening SR fields (Fig. S1).

In summary, the width and area of place fields within an environment showed the largest increase with distance to the nearest wall in the mismatch model, consistent with the experimental data, whereas the match and SR models showed similar trends of smaller magnitude. Across environments, the models diverged qualitatively: the SR model predicted decreasing place field area with increasing environment size, whereas the match and mismatch models predicted increases, consistent with the experimental data.

### Bayesian cue combination explains place field density

The precision of the visual likelihood further explains place field peak density in the mismatch model, via Bayesian cue combination (Eq. 14). When the agent is near a wall, it is much closer to that wall than to the opposite one. This means that although the visual information from the two walls may conflict, that from the nearer wall has much higher precision and therefore dominates the posterior estimate of location. In contrast, when the agent is far from both walls, the visual inputs are similarly imprecise, so the posterior estimate interpolates between them. As a result, the estimate changes more gradually with physical position. Under the simple assumption of approximately uniform tuning over the believed environment, this slower variation yields fewer place field peaks (Fig. 2b, Row 1, Fig. S2). Consequently, the mismatch model reproduces the experimental finding that the density of place field peaks decreases with distance from the wall (Fig. 2f; environment D; KW: *H* = 16.28, *p* = 0.001, MW: all *p <* 0.05 except 30–60 vs. 60–90 and 60–90 vs. 90–120 cm, n.s.; see also Fig. S4a, b for environments B and C.).

In contrast, the match and SR models failed to reproduce this pattern: because they do not distinguish between true and believed environments, the density of their place field peaks remains relatively constant regardless of distance from the wall (Fig. 2f; environment D; match, KW: *H* = 5.03, *p* = 0.17; SR, KW: *H* = 8.81, *p* = 0.03, MW: all comparisons n.s.).

A similar pattern holds across environments: place field peak density near a wall is lower in larger environments for the mismatch model (Fig. 2g; KW: *H* = 17.86, *p* = 0.0005, MW: all comparisons *p <* 0.05). This reflects the fact that the constant number of peaks in the mismatch model is distributed across a larger area (Fig. 2b, Row 3). The match and SR models do not reproduce this pattern (Fig. 2g; match, KW: *H* = 0.28, *p* = 0.96; SR, KW: *H* = 0.60, *p* = 0.90). Although the extent to which place field size depends on location within an environment and on environment size may vary with ideal-observer parameters, such as visual contrast (which lowers visual uncertainty) and self-motion noise (which increases transition uncertainty), the density pattern described here is qualitatively more robust.

### Confusability explains the scaling of active place cell counts

Using a fixed believed environment across different true environments, as in the mismatch model, also has measurable consequences for the overall level of place cell activity. In larger environments, a place cell is more likely to fire above threshold because more physical locations are confusable with the believed location to which the cell is tuned. The increase is sublinear with respect to area: once a neuron’s threshold is exceeded and a place field appears, additional confusable locations around existing fields do not create new fields. This yields a sublinear scaling of active place cell counts with environment area, a prediction characteristic of the mismatch model. In contrast, the match and SR models treat every location as independent, so the number of place cells grows linearly with area. Indeed, the mismatch model exhibited a sublinear trend with a substantially positive intercept that is consistent with experimental observations, whereas the match and SR models produced nearly linear trends and intercepts close to zero (Fig. 2h).

### Confusability explains place field shapes

In the previous sections, we showed that the mismatch model could reproduce key features of place cell activity across multiple environments by interpreting new sensory input using a fixed internal map. However, because the animals were similarly familiar with all environments in that experiment, we next considered a complementary setting in which animals were explicitly familiarized with a single environment and then placed into geometrically deformed versions of that environment that they had never experienced. This design more directly mirrors the assumptions of the mismatch model, where a fixed internal map is used to interpret unfamiliar input.

One such experiment comes from studies by O’Keefe and Burgess (1996), in which animals were placed into environments that appeared familiar but were actually stretched or compressed, creating a mismatch between the believed and true environments (Fig. 3a)^19^. Specifically, animals were familiarized in a vertically oriented rectangular environment and then tested in three deformed variants: a small square, a horizontally stretched rectangle, and a large square. For most cells that did not remap, the shape of the place field varied systematically with that of the environment, particularly along expanded axes. Specifically, in the horizontal rectangle and large square, fields either elongated along the direction of expansion or developed a second peak. Consistent with these findings, the mismatch model produced similar effects in its posterior place fields, with stretching and multi-peak patterns in expanded environments (Fig. 3b). These effects follow from the principles described above: posterior uncertainty is higher along longer axes (Fig. 2c) and mismatches between the believed and true environments can produce posteriors with multiple peaks (Fig. 1d), respectively.

**Figure 3.**
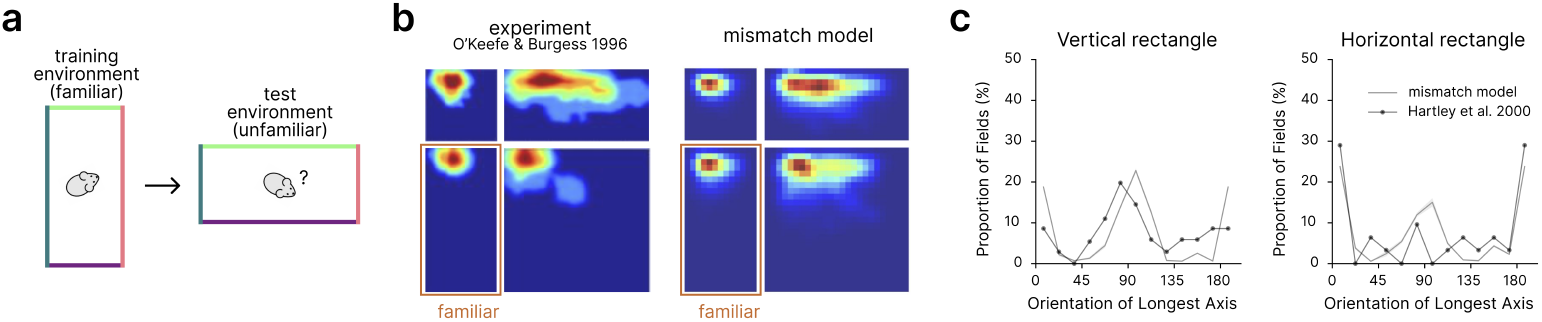
**a**. Schematic of an experiment designed to create a mismatch between the believed and true environments: the agent is trained in a familiar environment (vertical rectangle) and then tested in an unfamiliar, geometrically deformed version of that environment (here, horizontal rectangle). **b**. Example place fields from Ref. 19 (left) and from the mismatch model (right). In both cases, place fields stretch or develop additional peaks in the deformed environment. **c**. Distribution of place field orientations in the vertical and horizontal rectangles. The mismatch model (gray line) captures experimentally observed alignment of place field axes with the longest axis of the environment (black line; see Fig. S7a for full results)^35^.

**Figure 4.**
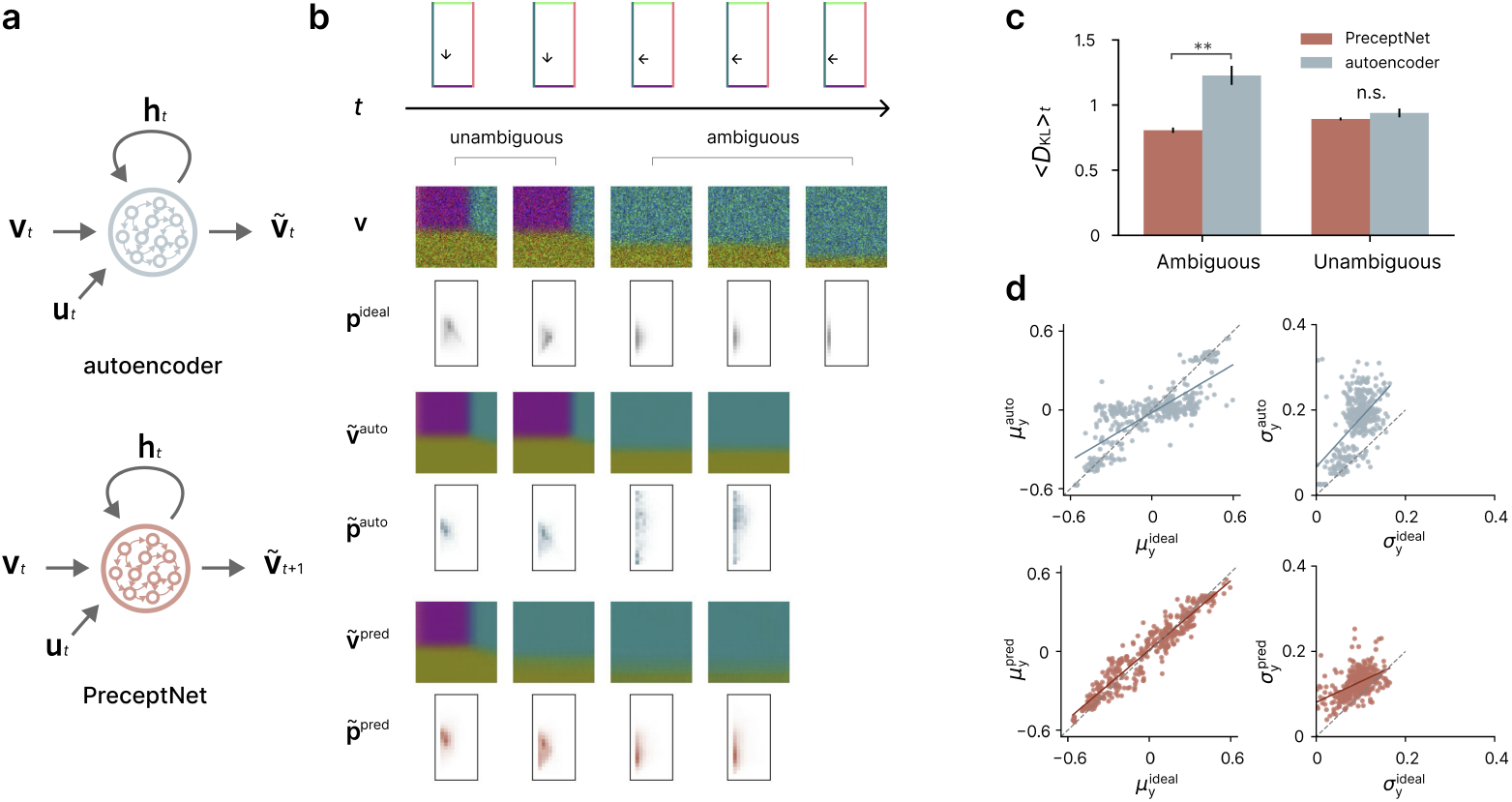
**a**. Architecture of the autoencoder (top) and predictive recurrent neural network (“Precept-Net”; bottom). Both networks receive the same egocentric inputs (visual input and self-motion), but are trained with different objectives: the autoencoder to reconstruct the current visual input, and PreceptNet to predict the next visual input. **b**. *Row 1*. Top-down view of the agent’s trajectory. The arrow indicates the current location and heading direction. *Row 2*. Noisy egocentric view of the 3D environment (**v**). *Row 3*. Posterior distribution over location calculated by the ideal observer. *Row 4*. Visual input reconstructed by the autoencoder. *Row 5*. Posterior distribution decoded from the autoencoder 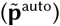. *Row 6*. Next visual input predicted by PreceptNet. *Row 7*. Posterior distribution decoded from PreceptNet 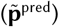 **c**. Average *D*_KL_ between decoded and ideal observer posterior distributions, shown separately for time steps with ambiguous and unambiguous poses. **d**. Scatter plots comparing the momentary mean (*µ*_y_) and standard deviation (*σ*_y_) of the decoded and ideal observer posteriors over the y coordinate at time steps with ambiguous poses.

A follow-up analysis by Hartley et al. further showed that the longest axis of the place fields consistently aligned with the longest axis of the deformed environment^35^. The mismatch model again reproduced this pattern, due to the same stretching mechanism described above (Fig. 3c and Fig. S7a).

Because the match and SR models assign independent place cell populations to different environments, they do not naturally account for neuron-specific transformations across environments.

### Network models of Bayesian representation learning

If a Bayesian belief over a latent state—here, the allocentric location—explains neural activity patterns, a key next question is how such neural representations can be learned solely from egocentric sensory inputs without external supervision. Previous work has shown that predictive learning can extract latent state representations from such inputs^13–17^. However, it remains unknown whether predictive learning also enables the encoding of Bayesian beliefs.

To test this possibility, we constructed a predictive recurrent neural network (“PreceptNet”) that learns to predict the next visual input from the history of visual and self-motion inputs (Fig. 4a, bottom). We designed the architecture so that its hidden layer receives the information necessary to recursively update beliefs about the latent state, as in Bayesian filtering (Fig. 6b): the current visual and self-motion inputs as well as its own previous value (which may correspond to the previous belief, i.e., the dynamic prior). Importantly, the network was not constrained to represent beliefs explicitly, nor was it given any direct supervisory signal about the latent state.

**Figure 5.**
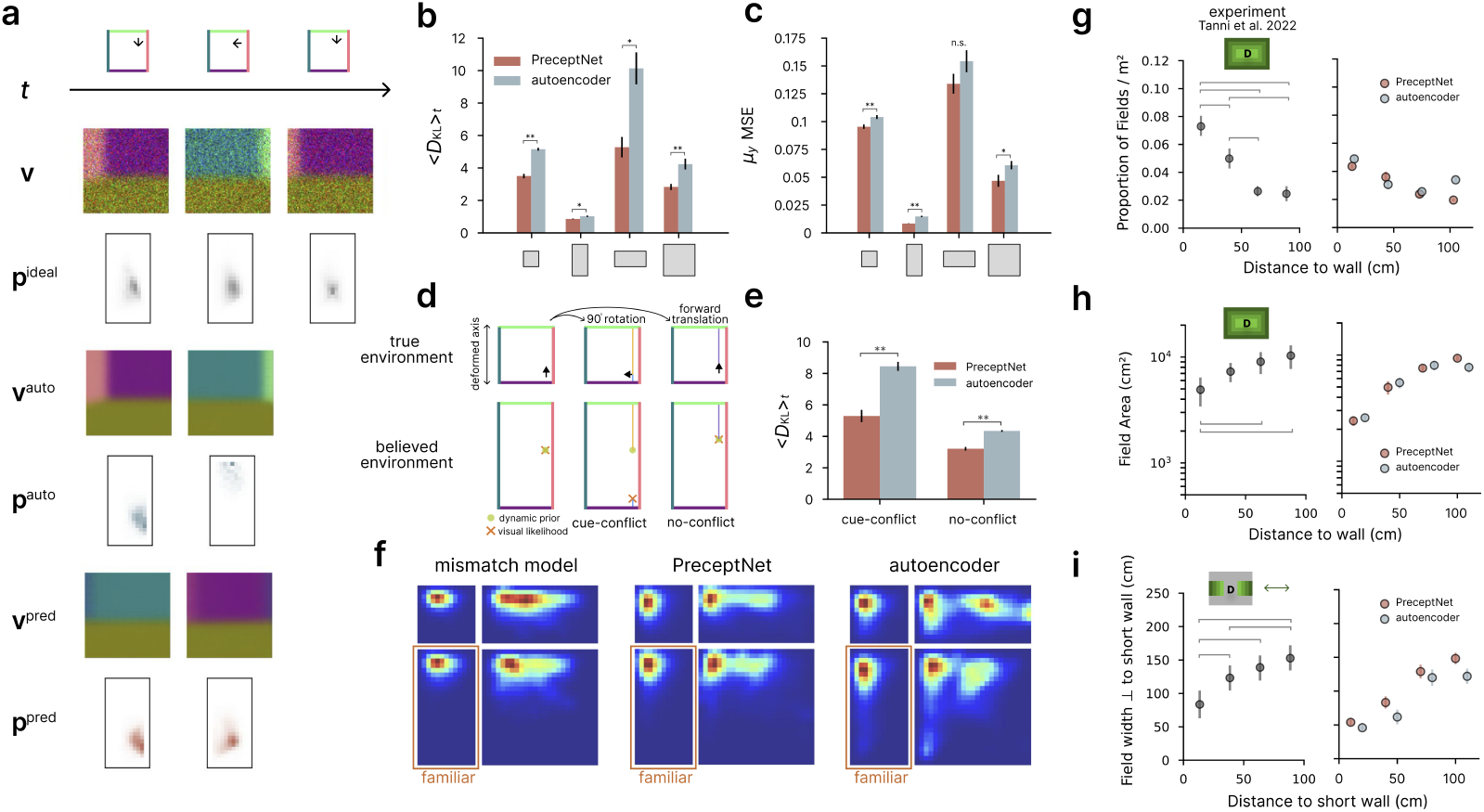
**a**. Example time steps in an unfamiliar environment (small square). Note the large divergence between 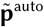 and 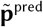 at the second time step, reflecting a cue conflict. See also panel d; conventions are the same as in Fig. 4b. **b**. Average *D*_KL_ between decoded and ideal observer posteriors across physical environments. **c**. MSE between decoded and ideal posterior means at each time step along the vertical axis across the same environments. **d**. Schematic illustration of cue-conflict and no-conflict time steps. In an unfamiliar environment, when the agent undergoes a 90° rotation, the dynamic prior (green dot) and visual likelihood (orange cross) suggest different positions along the deformed axis, leading to cue conflict. In contrast, during a forward translation (no-conflict), the dynamic prior and visual likelihood remain aligned, both indicating similar believed locations. **e**. Average *D*_KL_ between decoded and ideal observer posteriors for cue-conflict and no-conflict time steps. PreceptNet showed lower *D*_KL_ than the autoencoder in both cases, with a larger gap in cue-conflict time steps. **f**. Posterior place fields in deformed environments. PreceptNet yields spatially smooth, stretched fields; the autoencoder shows fragmented, irregular fields. **g–i** Comparison with place field statistics from Tanni et al. (2022). **g**. Field peak density as a function of distance to the wall (see also Fig. 2f). **h**. Field area as a function of distance to the wall (see also Fig. 2d). **i**. Field width along the axis orthogonal to the short wall (see also Fig. 2c).

**Figure 6.**
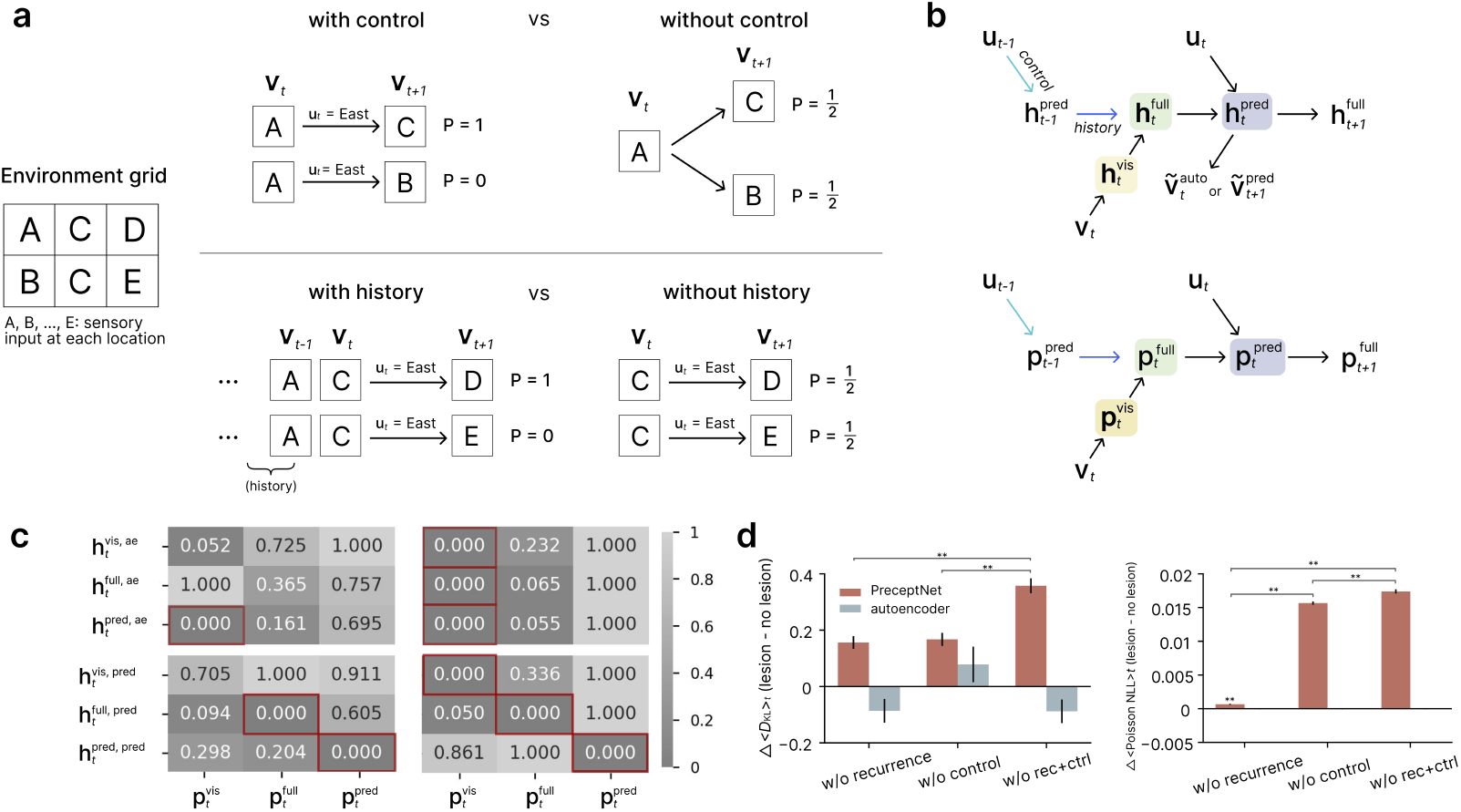
**a**. Toy environment illustrating how control (action) and history (recurrence) disambiguate the next observation. *Left*: Environment grid showing a 3×2 set of locations, each marked with the identity of the sensory input. Two middle locations provide the same sensory input (C), potentially introducing ambiguity. *Top*: Comparison of the predicted location distribution (predictive distribution) when the control information is available (left) or unavailable (right), when the current sensory input is A. *Bottom*: Comparison of the predictive distribution when the history information is available (left) or not (right), when the previous and current sensory inputs are A and C, respectively. See text for details. **b**. Information pathways in the network mapped onto the steps of Bayesian filtering (see also panel c). Cyan and blue arrows indicate connections ablated in simulated lesion studies targeting control and recurrent (history) inputs, respectively. **c**. Decoding heatmaps. Each cell shows normalized *D*_KL_ between the ideal observer posterior and the decoded posterior. *Left*: *D*_KL_ normalized within each column to range from 0 to 1; the red outline marks the hidden state best matching each target posterior. *Right*: *D*_KL_ normalized within each row to range from 0 to 1; the red outline marks the target posterior best matching each hidden state. **d**. Effect of input ablations on posterior decoding and predictive/autoencoding performance. *Left*: Change in *D*_KL_ with the ideal observer’s posterior comparing lesioned to intact networks. *Right*: Decrease in predictive performance, measured as the increase in Poisson NLL after lesioning. See Fig. S9a for comparison with the minimum and maximum possible *D*_KL_ and NLL values.

Specifically, the network first compresses the noisy egocentric visual input (**v**_*t*_) into a visual embedding 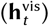. This embedding is combined with the previous predictive hidden state 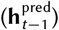 to form the full hidden state 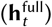, which is then combined with the self-motion input (**u**_*t*_) to yield the next predictive hidden state 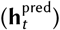. The resulting state is decoded to reconstruct the *next* visual input (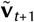; Fig. 6b, top).

As a control, we also trained another network (“autoencoder”) with the same structure and inputs as PreceptNet, but trained instead to reconstruct the *current* visual input (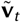; Fig. 4a, top). Because the autoencoder maps the current input directly to its reconstruction, we expected it to have limited capacity to integrate information across time, for example by incorporating the dynamic prior.

To examine whether PreceptNet’s network state, or that of the autoencoder, represents the ideal observer’s beliefs about location, we trained a separate decoder for each network. Each decoder mapped the network’s full hidden state 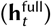 through a single fully connected (FC) layer followed by a softmax function, yielding a probability distribution over locations at the current time step (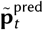 or 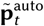), which we refer to as the decoded posterior. We optimized the FC layer to minimize the Kullback– Leibler divergence of the decoded posterior from the ideal observer posterior 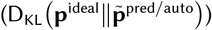. We then froze the decoder weights, applied the decoder to a held-out test set, and examined the resulting *D*_KL_.

### PreceptNet encodes posterior beliefs in a familiar environment, even under visual ambiguity

To test whether and how PreceptNet and the autoencoder encoded the ideal observer’s posterior beliefs about the latent state, we evaluated them in two types of environments: familiar and unfamiliar, as defined by O’Keefe and Burgess^19^. A familiar environment was one in which the network had been trained; all others were considered unfamiliar.

In a familiar environment, we found that both 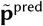 and 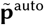 approximated the ideal observer’s posterior **p**^ideal^ reasonably well overall (Fig. 4b). However, at certain time steps, the decoded posteriors diverged more noticeably (Columns 3 and 4). To better understand these differences, we divided the agent’s poses (location and heading direction pairs) into two types: *ambiguous* poses, which share expected visual input with at least one other pose, and *unambiguous* poses, which have unique expected visual input. In a vertical rectangular environment, typical ambiguous poses included those facing the vertical wall, because visual inputs were identical along different positions along the vertical axis. If predictive learning encodes Bayesian beliefs, differences between the networks should be most apparent under ambiguous conditions. Indeed, separating time steps in the test set by pose ambiguity revealed a clear pattern (Fig. 4c). Critically, for ambiguous poses, PreceptNet showed significantly smaller *D*_KL_ values than the autoencoder (autoencoder: 1.23 ± 0.07, PreceptNet: 0.81 ± 0.02; paired *t*(4) = −4.89, *p* = 0.008), whereas for unambiguous poses, the difference did not reach significance (autoencoder: 0.94 ± 0.03, PreceptNet: 0.89 ± 0.01; *t*(4) = −1.10, *p* = 0.33).

In principle, there can be multiple reasons why the ideal observer’s posterior can be better decoded from one network than from another. For example, the decoded posterior may more closely match the mean estimate of location, or the uncertainty of the estimate. Thus, to understand the source of PreceptNet’s lower KL divergence, especially under ambiguous conditions, we compared the momentary mean and standard deviation of 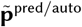 with those of **p**^ideal^ along the vertical axis, using only ambiguous inputs. We focused our analysis on this axis, as this is where the ambiguity primarily arose (analysis along the horizontal axis yielded qualitatively similar results; not shown). A clear difference between the two networks emerged from this comparison: when facing a long wall, many locations yield nearly identical visual inputs, and the autoencoder—relying only on the current input—cannot distinguish among them. Consequently, it often produced broad posteriors, with decoded means biased toward the center and inflated standard deviations. In contrast, PreceptNet more closely matched the ideal observer’s posterior both in terms of the mean and the standard deviation (Fig. 4d). Specifically, PreceptNet showed significantly lower mean squared error (MSE) between the momentary mean of its decoded posterior and that of **p**^ideal^ (autoencoder: 0.015 ± 0.001, PreceptNet: 0.008 ± 0.0001; paired t-test, *t*(4) = −7.35, *p* = 0.002). The same pattern held for the momentary standard deviation (autoencoder: 0.004 ± 0.0002, PreceptNet: 0.0002 ± 0.0001; paired t-test, *t*(4) = −6.60, *p* = 0.0002). These results suggest that by integrating information over time through its recurrent dynamics, PreceptNet more effectively captures both the location estimate and its uncertainty when visual input alone is insufficient.

### PreceptNet encodes posterior beliefs even in unfamiliar environments

Having shown that PreceptNet encodes posterior beliefs more accurately than the autoencoder in a familiar environment, we next asked whether this advantage generalizes to unfamiliar environments. We froze all network parameters, including those of the posterior decoder, after training in the familiar vertical rectangle environment and evaluated their performance in unfamiliar environments, following the procedure used for animals in the same experiment^19^. This procedure mirrors that used for the mismatch model discussed earlier, in which the believed environment remains fixed even when the true environment changes.

Across all three unfamiliar environments tested, PreceptNet’s decoded posterior matched the ideal observer’s posterior significantly better than the autoencoder’s, with consistently lower *D*_KL_ values (Fig. 5b). In addition, the momentary posterior mean along the vertical axis decoded from PreceptNet showed lower MSE than that from the autoencoder across environments; this difference was significant in two of the three environments (small and large squares), with a comparable trend in the horizontal rectangle (Fig. 5c).

To better understand these differences, we focused on *cue-conflict* time steps, when visual and self-motion signals provide conflicting information about the agent’s location. Such steps occur frequently in unfamiliar environments and likely contribute substantially to the performance gap between networks. By contrast, *no-conflict* steps are those in which the cues agree. We examined two types of cue-conflict: 90° rotations (Fig. 5d) and 180° rotations (Fig. S8). Both produce a sharp divergence between the believed location (in the familiar vertical rectangle) and the true location (in the deformed environment). Because the agent rotates in place, the dynamic prior, updated only through self-motion, retains its peak at a similar location, whereas the peak of the visual likelihood shifts abruptly to the position in the believed environment consistent with the newly faced wall. Reconciling this discrepancy requires effective use of the dynamic prior.

Indeed, in cue-conflict steps, PreceptNet’s decoded posterior shifted smoothly, integrating information from both sources and closely matching the ideal observer’s beliefs. In contrast, the autoencoder’s posterior mean jumped abruptly toward the location implied by vision alone (Fig. 5a), yielding a significantly larger average *D*_KL_ from the ideal observer’s posterior (Fig. 5e; autoencoder: 8.45 ± 0.28; PreceptNet: 5.29 ± 0.39; paired t-test, t(4) = -11.15, *p* = 0.0003). A similar difference in *D*_KL_ was observed in no-conflict steps, though it was smaller (autoencoder: 4.29 ± 0.03, PreceptNet: 3.19 ± 0.11; paired t-test, t(4) = -13.41, *p* = 0.0002). The magnitude of this difference was significantly greater in cue-conflict than in no-conflict steps (cue-conflict steps: 3.16 ± 0.28, no-conflict steps: 1.10 ± 0.08; paired t-test, t(4) = 6.97, *p* = 0.002, see also Fig. S8 for cue-conflict during 180° rotations).

Together, these results suggest that PreceptNet benefits from more effective use of the dynamic prior in unfamiliar environments. By contrast, the autoencoder’s representations appear dominated by instantaneous visual likelihoods. This motivates the next question: why prediction is necessary for making effective use of the dynamic prior.

### Prediction drives encoding of posterior beliefs

To illustrate why prediction enables effective use of a dynamic prior, we begin with a simple example. Here, sensory inputs are deterministic: each grid location yields a fixed visual observation denoted by a letter (e.g., “A”, “B”; Fig. 6a). Prediction ambiguity can be reduced by incorporating control information: knowing the agent’s action (e.g., moving east) helps disambiguate the expected next location (Fig. 6a, top). However, because different locations can yield the same observation, uncertainty can remain even when both the current observation and action are known. In such cases, identifying the correct next location (and hence next observation) also depends on the agent’s prior trajectory— information that must be carried over time through recurrence (Fig. 6a, bottom). These examples illustrate that satisfying the predictive objective requires not only perceptual input, but also control and history. In contrast, an autoencoding objective in this example does not: a perfect autoencoder’s internal state need not distinguish between locations with an identical visual input (“C”).

In probabilistic terms, it suffices for the autoencoder to encode the posterior based solely on visual input (the vision-only posterior, **p**^vis^ := P(**s**|**v**)) in this example, since further disambiguation of the state is unnecessary when the state maps to a visual input deterministically. Consistent with this intuition, even in the original stochastic task, all of the autoencoder’s hidden states (**h**^vis^, **h**^full^ and **h**^pred^) are found to encode **p**^vis^ best (Fig. 6c right, top three rows).

In contrast, even when visual inputs are deterministic, PreceptNet must make use of the history and control inputs. That is, its state after receiving the history input (**h**^full^) must correspond to the full posterior, **p**^full^ := P(**s**|**v**_0:*t*_, **u**_0:*t*−1_), and that after receiving the control input (**h**^pred^), to the predictive distribution, **p**^pred^ := P(**s**|**v**_0:*t*_, **u**_0:*t*_). Analytical derivations for large shallow networks show that the predictive objective leads to **h**^full^ that encodes the full posterior, while the autoencoding objective yields representations that encode the vision-only posterior (Supplementary Information). Numerical results confirm the same pattern in our deep convolutional networks (Fig. 6c right, bottom three rows).

Having established that prediction gives rise to posterior-like representations, we numerically confirmed the contribution of recurrent and control inputs. For this, we ablated each of the recurrent and/or the control inputs (Fig. 6b, blue and cyan arrows, respectively), corresponding to the history and self-motion inputs, respectively, by setting those input vectors to zero during training.

For PreceptNet, removing either recurrence or control significantly worsened the decoded beliefs and jointly removing both produced the largest degradation (Fig. 6d left; Δ*D*_KL_, w/o recurrence: 0.16 ± 0.02, w/o control: 0.17 ± 0.02, w/o both: 0.36 ± 0.03). Importantly, every disruption of posterior representation was accompanied by a decline in accuracy in predicting the next sensory input, reflected in an increase in target loss (PoissonNLL), with the same general pattern across lesions (Fig. 6d right; ΔPoissonNLL, w/o recurrence, w/o control: 0.016, 0.0007, w/o both: 0.017). This pattern is consistent with the idea that the predictive objective encourages PreceptNet to encode the posterior better.

In contrast, the autoencoder showed little to no change in either *D*_KL_ or target loss, indicating that it did not effectively rely on recurrence or control to optimize its objective (Δ*D*_KL_, w/o recurrence: -0.09 ± 0.04, w/o control: 0.08 ± 0.06, w/o both: -0.09 ± 0.04; ΔPoissonNLL, w/o recurrence: − 4.9 *×* 10^−5^, w/o control: −5.6 *×* 10^−5^, w/o both: −4.8 *×* 10^−5^). Notably, recurrence was especially critical for posterior encoding in ambiguous poses: removing recurrence abolished PreceptNet’s advantage over the autoencoder (Fig. S9b, top left vs. bottom left), and further removing control input increased PreceptNet’s disadvantage (Fig. S9b, bottom left vs. bottom right). Together, these results support the conclusion that the predictive objective promotes posterior encoding by compelling the network to exploit recurrent and control inputs.

### PreceptNet’s activity resembles place cell activity

So far, we have shown that PreceptNet represents the ideal observer’s beliefs (Fig. 4 and Fig. 5a–f), which in turn resemble place cell activity (Figs. 2 and 3). We next asked whether PreceptNet’s activity directly resembles place cell activity. To test this, we computed posterior place fields by averaging the decoded posterior distributions over time, using the same method as for the ideal observer models.

In simulations of the O’Keefe and Burgess study^19^, we found that PreceptNet produced place fields that systematically stretched along the axis of environmental expansion, closely resembling both biological place cells and the mismatch model (Fig. 5f and Fig. 3b; see also Fig. S7). In contrast, the autoencoder yielded less smooth place fields, often with irregular shapes or multiple peaks. These differences are consistent with the autoencoder’s sensitivity to momentary visual input, particularly under cue conflict, where visual inputs from the same location in the familiar environment can occur at different physical locations in the unfamiliar environment (Fig. 5d). However, these qualitative differences alone were not sufficient to conclude whether PreceptNet or autoencoder resembled place cells better. Indeed, both networks closely matched the distribution of peak orientations observed in place cells (Fig. S7b).

Therefore, we turned back to the Tanni et al. experiment for quantitative comparison of the networks^24^. A major difference between the two networks emerged in the central region of the environment: in the autoencoder, place fields were often fragmented, resulting in a higher density of peaks, smaller areas, and reduced widths along the axis orthogonal to the short wall, patterns that are inconsistent with experimental findings.

In contrast, PreceptNet reproduced the key spatial trends seen in biological data. Specifically, peak density decreased (Fig. 5g; env. D; PreceptNet, KW: *H* = 17.58, *p* = 0.0005, MW: all *p <* 0.05; autoencoder, KW: *H* = 16.00, *p* = 0.001, MW: all *p <* 0.05, except 30–60 vs. 60–90, 90–120 cm, n.s.), and both field area and field width increased with distance from the wall (Fig. 5h; PreceptNet, KW: *H* = 17.86, *p* = 0.0004, MW: all *p <* 0.05; autoencoder, KW: *H* = 16.14, *p* = 0.001, MW: all *p <* 0.05 except 60–90 vs. 90–120 cm, n.s.; Fig. 5i; PreceptNet, KW: *H* = 17.33, *p* = 0.0006, MW: all *p <* 0.05; autoencoder, KW: *H* = 16.10, *p* = 0.001, MW: all *p <* 0.05, except 60–90 vs. 90–120 cm, n.s.; see also Fig. S10).

Taken together, these results indicate that PreceptNet’s activity not only mirrors the mismatch model’s posterior beliefs but also captures key spatial features of biological place cell activity.

## Discussion

Uncertainty is unavoidable in perception and action: sensory evidence is noisy and ambiguous, and actions have stochastic consequences. For robust behavior, internal representations must therefore encode not only point estimates of latent variables, such as allocentric location, but also their uncertainty. It has long been noted that humans and other animals estimate and use such uncertainty in their behavior in an approximately optimal fashion, i.e., following Bayesian principles^36^. However, it has remained unresolved how the brain represents Bayesian beliefs and, critically, how such representations could be learned from the sensory information available to the animal. Here we address both questions by showing that hippocampal place cells encode posterior-like representations over allocentric location, and that such representations can be learned from egocentric sensory input streams alone, through predictive learning.

Specifically, we first compared three computational models of spatial coding: the widely used SR, which captures expected future occupancy based on transition dynamics alone^10^, and two Bayesian ideal observer models, which incorporate both transition and perceptual uncertainty assuming either a match or a mismatch between the agent’s internal map and the physical environment. Surprisingly, systematic variations in place field properties (such as density and size) within and across environments were best explained by corresponding variations in the mismatch model’s beliefs. These, in turn, were determined by optics, which set the precision of egocentric visual input, and by Bayesian cue combination. The SR failed to account for these variations because it ignores systematic changes in sensory ambiguity.

We next asked whether such spatial representations, encoding both state estimates and their associated uncertainty as in the ideal observer model, could be learned from experience. To address this, we trained an RNN, termed PreceptNet, to predict future egocentric visual input based on the history of visual and self-motion inputs. Remarkably, the network developed internal states that closely resembled the posterior distributions of a Bayesian ideal observer. These belief-like representations gave rise to place cell-like activity that generalized across unfamiliar environments and reproduced the key spatial trends observed in the brain. By contrast, an otherwise identical network trained with an autoencoding objective, the autoencoder, did not form comparably faithful posterior codes or place fields. This supports the role of the predictive objective in driving the emergence of posterior-like neural representations.

The difference between PreceptNet and the autoencoder emerged most clearly in ambiguous states, underscoring the central role of perceptual uncertainty. Although real-world sensory inputs at different poses are rarely identical, they are often highly similar and therefore ambiguous. Such near-degeneracies arise generically in environments with repeating structure, for example along paths lined with uniform walls or visually similar trees. In these regimes, predictive learning confers an advantage.

### Comparison with prior works

Relative to one-step SR^10,18^, our framework approximately constitutes a superset: one-step SR captures self-motion–based prediction, and adding visual evidence as in our framework yields the full posterior (Fig. 1a and One-step SR model). For multi-step SR, the relationship is complementary: discounted future occupancy captures place cell activity patterns that our framework does not (e.g., splitter cells^37,38^), but it cannot explain within- and across-environment trends (Figs. 2 and 3). It is straightforward to extend our model to multiple steps, in which a place cell would represent the expected frequency of visits while accounting for state uncertainty^39^.

While previous studies of cognitive maps have used predictive objectives to extract spatial latent variables^13–15,17^, our study is unique in that we treat the latent code as a probability distribution, not merely a point estimate. By directly comparing network states against an ideal observer posterior, we show that predictive learning yields representations that track both estimated location and its associated uncertainty.

Other studies have demonstrated the emergence of probabilistic representations in neural networks^40–42^. However, these approaches rely on supervision with the ground-truth posterior^41^, target outputs^42^, or external reinforcement^40^. Such forms of supervision or reinforcement are rare in the real world, making them implausible mechanisms for learning the association between streams of complex sensory input and high-dimensional latent states, as required in natural tasks such as navigation.

More broadly in the machine learning community, prior work has shown that Bayesian filtering–like computations can be implemented in recurrent neural networks^43,44^. However, these approaches did not directly test whether internal states correspond to Bayesian ideal-observer posteriors. In contrast, our model is trained only with a self-supervised predictive objective, yet induces hidden representations from which individual stages of Bayesian filtering—likelihood, posterior, and predictive distribution—can be decoded and quantitatively matched to an ideal observer on a moment-by-moment basis. This framework further provides a direct link to hippocampal spatial representations, showing how predictive learning under perceptual ambiguity can give rise to place cell–like activity without explicit probabilistic supervision or engineered inference rules.

### Biological predictions

Hippocampal connectivity exhibits features well suited for encoding Bayesian posterior beliefs. It receives convergent egocentric self-motion information (e.g., from parietal and vestibular systems^45–47^), and egocentric visual information (e.g., from entorhinal and retrosplenial cortices^48–52^), providing the ingredients required for Bayesian updates. Moreover, its rich recurrent circuitry, particularly in CA3, may provide the dynamic prior in our framework.

Our ideal observer model makes quantitative predictions for place fields when sensory (visual or self-motion) or recurrent inputs are manipulated. Increasing visual reliability (e.g., adding a landmark) should cause place fields to cluster, reflecting stronger anchoring by precise visual likelihoods, and to become narrower, reflecting reduced uncertainty (as in Fig. 2c–e)^31,53^. Decreasing self-motion reliability (e.g., via vestibular or locomotor disruption) should broaden the dynamic prior, producing larger fields with lower spatial information^54^. Introducing conflicts between vision and self-motion should cause field locations to interpolate between the cues, especially for small conflicts (as in Fig. S2)^20,55^. Weakening recurrence should destabilize fields and make them more dominated by sensory cues, consistent with reports that CA3 disruption yields place fields with reduced sharpness, along with impaired spatial recall^56^.

Our neural network results further suggest that place field formation depends on the predictability of upcoming sensory input. This aligns with experiments showing that hippocampal cells lose selectivity when sensory sequences are made unpredictable^57^. It also predicts that place fields may fail to form in virtual reality experiments if the predictability of movement outcomes is degraded, or when movement is prevented and only shuffled snapshots of a place are presented. This prediction would not follow if place fields were merely compressed memory representations, independent of action or temporal order (as in our autoencoder and in Ref. 58 without extension).

### Applications and broader impact

Notably, the mismatch model—the Bayesian model assuming a mismatch between the internal map and the physical environment—provided the best account of biological place fields. In our implementation, this internal map was fixed at an average geometry, approximating Bayesian averaging. While this choice might seem extreme, a relatively fixed internal map could be sensible when the animal is uncertain about environmental geometry, which is likely when it experiences multiple large environments with non-negligible visual and self-motion uncertainty. The precise representation of such map uncertainty remains an intriguing topic for future research. The superior fit of this model, relative to separate veridical internal maps, offers a rudimentary example of inference about the structure of the internal map—in addition to the subjective belief about location within it—from hippocampal activity. This suggests a direction for future work: decoding the detailed structure of the internal map from population activity^59^.

Beyond neuroscience, our predictive learning framework offers practical advantages for robotics and artificial intelligence. In robotics, and especially in simultaneous localization and mapping (SLAM), predictive models of this kind could approximate belief-like inference without explicitly engineered probabilistic algorithms. In artificial intelligence, reinforcement learning and model-based planning often rely on deterministic internal models that fail under ambiguity or partial observability. Our results suggest that predictive learning may suffice for the emergence of uncertainty-aware codes, enabling AI systems to integrate conflicting evidence and generalize more effectively to novel environments. While predictive learning has previously been used for pretraining^60^, our framework makes explicit the correspondence between internal representations and each stage of Bayesian filtering, providing interpretability.

More broadly, while spatial navigation offers a domain where the mapping between latent variables, observations, and their neural correlates is particularly clear, the same computational challenge— inferring hidden states from noisy, ambiguous input—extends to decision-making, active sensing, and other cognitive domains. Across these domains, a key ingredient for robustness is not only estimating the most likely state but also maintaining a representation of uncertainty^1^.

### Conclusion

Animals often show a remarkable ability to make near-optimal decisions under uncertainty, yet the neural underpinnings and learning mechanisms behind such competence have remained unclear. Here we address both: we show that hippocampal activity is consistent with belief-like representations, and that such codes can emerge through predictive learning. Thus, our framework bridges Bayesian inference over time with neural network training, and offers a generalizable principle: the brain can form belief-like codes from perceptual inputs and control signals under uncertainty.

## Methods

### Simulated environment and state space

All environments were discretized into uniform grids. For simulations replicating Tanni et al. (2022), which used large environments, we used a spatial resolution of 10 cm to avoid excessive demand on memory and compute; for simulations replicating O’Keefe and Burgess (1996), with modest environment sizes, we used a finer resolution of 5 cm. At each grid location, the agent could assume one of four discrete headings: 0°, 90°, 180°, or 270°. This yielded a state space consisting of all combinations of discrete locations and orientations, i.e. *N*_loc_ *×* 4 total states, where *N*_loc_ is the number of grid locations in the environment.

### Bayesian ideal observer model

We use the Bayesian ideal observer of navigation (BION) introduced in Ref. 11; here we summarize the relevant aspects. BION infers the agent’s allocentric state (location and heading direction) based on a sequence of egocentric visual observations and self-motion inputs. This inference is implemented via recursive Bayesian filtering, where the posterior distribution over the current state is updated at each time step by combining predictions from the previous state with new sensory evidence.

Let the agent’s allocentric state at time step *t* be **s**_*t*_, and let the noisy egocentric visual input be **v**_*t*_ . The observer also receives a self-motion signal **u**_*t*_, indicating intended translation and rotation. The posterior distribution **p**_*t*_ := *P*(**s**_*t*_ | **u**_0:*t*_, **v**_0:*t*_) is computed by combining three components:

- **visual likelihood** *P*(**v**_*t*_ | **s**_*t*_): the probability of the current visual input given the agent’s state;
- **transition model** *P*(**s**_*t*_|**u**_*t*_, **s**_*t*−1_): the probability of transitioning to the current state given the previous state and self-motion input;
- **dynamic prior** *P*(**s**_*t*−1_ | **u**_0:*t*−1_, **v**_0:*t*−1_): the posterior from the previous time step (**p**_*t*−1_).

With them, the posterior is updated recursively with Bayesian filtering. First, in the *prediction step*, the dynamic prior is combined with the transition model to yield the one-step predictive distribution, propagating the belief forward in time:

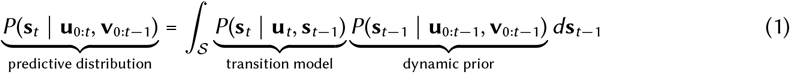

Then, in the *measurement step*, the predictive distribution is combined with the visual likelihood with Bayes rule to yield the posterior, incorporating the new visual observation to refine the belief:

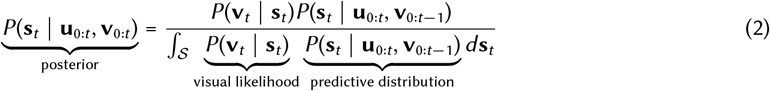

The ideal observer thus maintains a posterior **p**_*t*_ over the full allocentric state space, incorporating uncertainty in both transition and perception.

Concretely, the transition model was a Gamma distribution with mean one spatial bin and coefficient of variation 0.2. The visual likelihood was a Poisson distribution with rate given by the expected egocentric visual input; its noisiness was controlled by a contrast parameter set to 0.3. Full details are available in Ref. 11.

To simulate trajectories, we predefined a list of random goal states and let the agent sequentially navigate between them. At each step, the agent selects a direction toward the current goal and samples an intended translation and rotation. To simulate between-subject variability, five independent trajectories (random seeds) were generated for the Tanni et al. experiment (five animals) and the O’Keefe and Burgess experiment (six animals).

### Posterior place field acquisition

To compute the place field of a given neuron *i*, we assume its firing rate *r*_*i*_ reflects the posterior probability of being at a particular *believed* spatial location 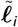, marginalizing over heading directions. Specifically, the neuron’s firing rate at each time step *t* was computed as:

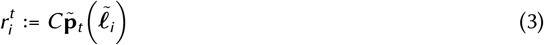

where *C* is a scaling constant chosen to match the mean firing rate observed in the experimental data. We then compute the spatial firing rate map of this neuron by averaging its activity across all time steps at which the agent was at a given *true* allocentric location ***ℓ***:

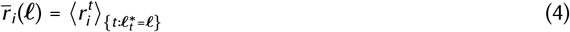

This results in firing rates defined over the layout of the *true* environment, simulating what would be measured experimentally.

We used a number of time steps proportional to the area of each environment: 3,000 time steps for environment A, 6,000 for B, 12,000 for C, and 24,000 for D in the Tanni et al. experiment; and 5,000, 10,000, 10,000, and 20,000 time steps respectively for the small square, vertical rectangle, horizontal rectangle, and large square in the O’Keefe and Burgess experiment. Place fields were then identified as continuous spatial regions where the neuron’s activity exceeded 50% of its peak value and the peak firing rate exceeded 2 Hz, consistent with thresholding procedures used in experimental studies^35,61^ (for a lower threshold, see Fig. S6).

### Successor representation model

The SR associates each location in the environment with a predictive vector encoding the expected future occupancy of all other states, discounted over time^9,10^. Formally, the SR matrix **M** is defined as:

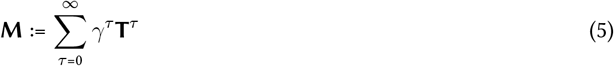

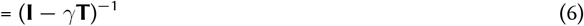

where *γ* is the temporal discount factor, and

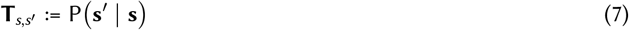

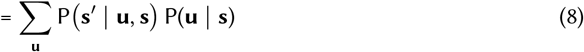

is the transition matrix. Here, the policy, P(**u**|**s**), is the probability of taking action **u** in state **s**, and the transition probability, P(**s***′*|**u**,**s**), is the probability of transitioning to state **s***′* given **u** and **s**. To estimate the SR, we used the same simulated trajectories as in the ideal observer model, generated by an agent navigating toward randomly selected goal states. The transition probabilities were then estimated empirically from these trajectories.

Each row of **M** represents the expected discounted future occupancy of all states starting from the corresponding state. To generate SR-based place fields, each neuron was assigned to a target state, and its activity across space was given by the corresponding column of **M**, reflecting the expected discounted visitation of that state from all other starting states. To ensure comparability with the ideal observer model, we used the same number of simulation time steps, environments, and thresholding criteria for defining place fields.

We tested values of *γ* ranging from 0.95 to 0.99 and found that the resulting SR matrices produced qualitatively similar results; we therefore used *γ* = 0.95 for all analyses.

### Place field orientation

To quantitatively compare the activity patterns of the models with those of place cells in deformed environments, we replicated the analyses in Refs. 19 and 35. To match the composition of the experimental data set, which included 28 units (21 recorded after pre-training in a vertical rectangle and 7 in a horizontal rectangle), we trained 325 place cell-like units (matching the number of spatial bins) in the vertical rectangle environment, randomly selected 240 of these units for the vertical rectangle condition, and generated 80 additional units for the horizontal rectangle condition by rotating the activity of a separate random subset by 90°.

To quantify place field orientation, we measured the axis of greatest spatial extent for each field, rounded the orientation to the nearest 7.5°, and binned the values into intervals of 15° to obtain an orientation distribution, as in Ref. 35.

### PreceptNet architecture and training

The PreceptNet architecture parallels the computational steps of Bayesian filtering and consists of four modules: a visual encoder, a recurrent integration module, a control-based prediction module, and a visual decoder. Together, these modules enable the network to integrate visual input with self-motion signals to predict future sensory input.

#### Visual encoder

Egocentric visual input **v**_*t*_ was an RGB image represented as a *C × H × W* tensor; in our implementation, we used 3 *×* 80 *×* 80. Observation noise in the visual channel was modeled as Poisson, consistent with the Poisson negative log-likelihood loss used for training. The images were processed through three convolutional layers with ReLU activations and 2 *×* 2 max pooling. The layers had channel transitions Conv1 (3 *×* 3 kernel, 3 → 32 channels), Conv2 (3 *×* 3, 32 → 16), and Conv3 (3 *×* 3, 16 → 8), each with stride 1 and padding 1. The resulting feature maps were flattened into a vector of size 800 and transformed by two fully connected (FC) layers (800 → 256 → 128) to produce a 128-dimensional visual embedding 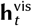.

#### Recurrent integration module

To integrate current visual evidence with accumulated past information, the visual embedding 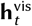 was concatenated with the predictive hidden state from the previous time step 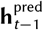 (128 dimensions). The resulting 256-dimensional vector was passed through FC layers (256 → 192 → 128) to yield the full hidden state 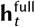. This module aggregates current sensory input with the network’s temporal memory, serving as an analogue to the Bayesian measurement update.

#### Control-based prediction module

To incorporate self-motion information, the full hidden state 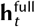 was concatenated with a 6-dimensional egocentric control signal **u**_*t*_ (2 dimensions encoding movement versus stop and 4 encoding rectified sine and cosine of heading change). This combined vector was processed by two FC layers (134 → 131 → 128) to produce the updated predictive hidden state 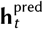. This module implements a control-dependent update analogous to the Bayesian prediction step, propagating internal state estimates according to intended motion.

#### Visual decoder

The predictive hidden state 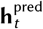 was decoded to generate the predicted visual input 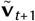 First, it was transformed through FC layers (128 → 256 → 800), reshaped into a feature map of size 8 *×* 10 *×* 10, and processed by three transposed convolutional layers: Deconv1 (3 *×* 3, 8 → 16 channels), Deconv2 (3 *×* 3, 16 → 32), and Deconv3 (3 *×* 3, 32 → 3), all with stride 2, padding 1, and output padding 1. ReLU activations were used throughout, with a final sigmoid to normalize pixel intensities.

#### Training

The network was trained to minimize the Poisson negative log-likelihood between the predicted and ground-truth future images (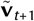 and 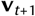), consistent with the Poisson observation noise. All weights were initialized using PyTorch’s default initialization (PyTorch v2.0.1) and optimized with Adam (learning rate 0.001) for 40 epochs with a batch size of 1. For simulations corresponding to Ref. 19, training was performed only in the vertical rectangular environment using 30,000 time steps. For simulations corresponding to Ref. 24, the number of training time steps scaled with environment size (15,000, 25,000, 35,000, and 45,000 time steps for environments A–D). The network was never directly provided with supervisory signals about the latent state.

### Autoencoder architecture and training

The architecture, input, and training configuration of the autoencoder were identical to those of PreceptNet; the only difference was in the training objective: While PreceptNet was trained to predict the next visual input, the autoencoder was trained to reconstruct the visual input at the current time step, **v**_*t*_ .

### Decoding posterior distribution

Regardless of whether the believed environment 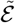 is the same as or different from the true environment ℰ^∗^, the ideal observer posterior 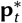 at time step *t* is of length 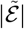, the number of spatial bins in the believed environment. The decoded posterior belief 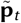 is of the same length, and is decoded from the hidden state **h**_*t*_ with weights 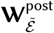 of dimensions 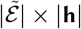, where |**h**| is the length of **h**_*t*_ :

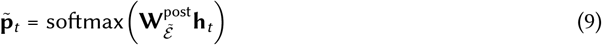

The belief of being at a particular state 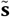 is the 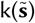-th element of this decoded vector (where k(·) is the index of a state), denoted simply as 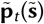. Note that **h** can denote any of **h**^vis^, **h**^pred^, or **h**^full^ (as in Fig. 6c) all of which have the same dimensionality. Likewise, the decoding target can be **p**^vis^, **p**^pred^, or **p**^full^.

## Acknowledgments

This work was supported by the National Research Foundation of Korea (NRF), funded by the Ministry of Science and ICT (RS-2025-16066185 and RS-2026-25492934), and by the International Joint Research Project at KAIST, also funded by the Ministry of Science and ICT of the Republic of Korea.

## Author Contributions

Y.H.R.K. conceived the study. Y.K. and Y.H.R.K. designed the simulations and analyses. Y.K. performed the simulations and analyses. Y.K. and Y.H.R.K. interpreted the results and wrote the manuscript.

## Supplementary Information

### Why SR place fields are smaller in larger environments

The SR is defined as the sum of discounted transition distributions over time:

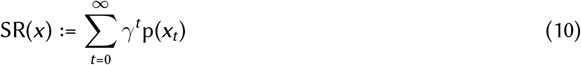

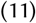

where *x* denotes location, *t* is the time step, p(*x*_*t*_) is the transition probability, and *γ* is the discount factor.

In an unbounded 1D environment, the transition probability from a state *s* can be approximated as a Gaussian distribution when the agent undergoes a random walk:

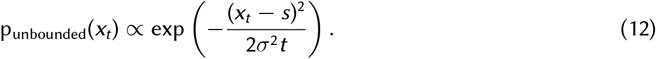

where *σ* reflects the spread determined by the agent’s movement variability.

In a bounded environment, the agent cannot move beyond the walls. We model this constraint by treating the walls as reflecting boundaries in the corresponding diffusion equation. The solution is obtained with the *method of images*, which introduces virtual reflected Gaussians placed at mirrored positions outside the environment:

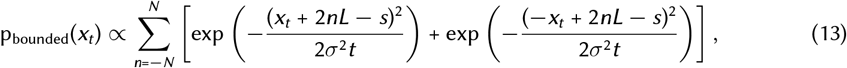

where *L* is the environment length and *N* is chosen to ensure numerical convergence.

When the environment size *L* is smaller, the reflected components are located closer to the encoding location *s*, leading to stronger overlap with the primary Gaussian and effectively redistributing probability mass back into the interior near the boundaries. This increased overlap elevates the SR values more broadly across the space. In contrast, in larger environments, the reflected components are positioned farther from the encoding location, and their influence becomes weaker, resulting in a more localized SR profile.

To numerically illustrate the effect of reflecting boundaries, we simulated p_unbounded_(*x*_*t*_), p_bounded_(*x*_*t*_), and their corresponding successor representations SR_unbounded_(*x*) and SR_bounded_(*x*) for a neuron encoding the center position *s* = *L/*2 in 1D environments of lengths *L* = 150 cm and *L* = 200 cm. We used a movement variability of *σ* = 15 cm, a discount factor of *γ* = 0.99, and truncated the SR sum at *t* = 30 time steps. The SR place field width was defined as the spatial extent over which each SR(*x*) exceeds 50% of its peak, as in the main analyses.

Fig. S1 illustrates how reflecting boundaries affect both the transition distribution over position after 10 time steps, p(*x*_10_), and the successor representation SR(*x*) in environments of both sizes. Fig. S1a,b show p(*x*_10_) with and without reflecting boundaries: in the small environment, reflections broaden the distribution, whereas in the larger environment the distribution is only weakly affected. Fig. S1 c,d show the corresponding successor representations SR(*x*), obtained by summing discounted distributions. The broader transition distribution p_bounded_(*x*_*t*_) in the small environment accumulates into a substantially wider SR place field, whereas in the larger environment, the SR remains tightly localized and close to the unbounded case (i.e., SR_unbounded_(*x*), which has identical shape across environments).

These simulations support the analytical intuition: boundaries have a stronger impact in smaller environments, leading to broader SR place fields.

**Figure S1.**
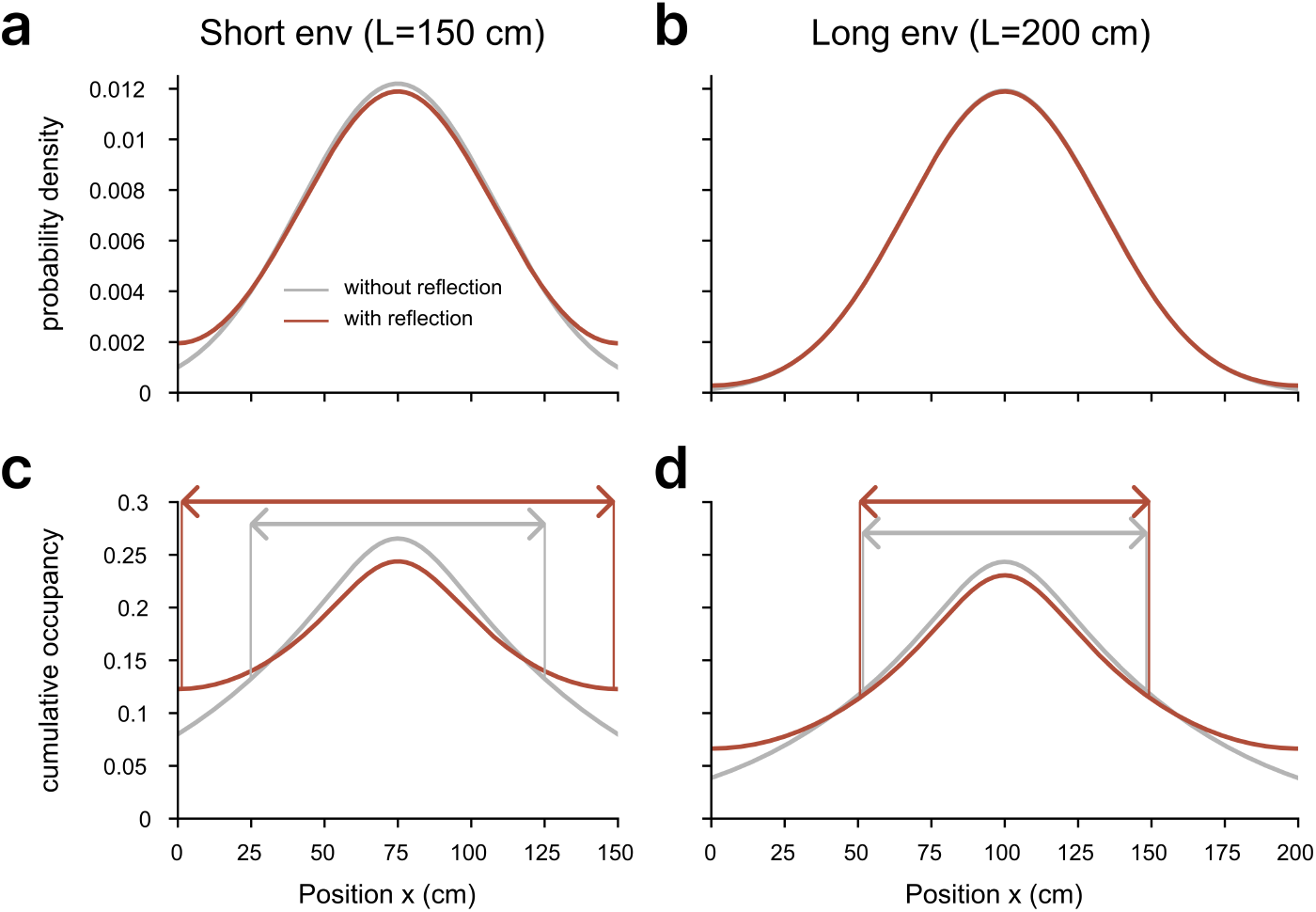
Effect of reflecting boundaries on position distributions and SRs in 1D environments. **a, b** Transition probabilities after 10 time steps, p_unbounded_(*x*_10_) (gray; without reflection) and p_bounded_(*x*_10_) (red; with reflection), in small (left, 150 cm length) and large (right, 200 cm length) environments. In the small environment (a), reflections produce a broader distribution by increasing probability density near the boundaries, leading to a flatter overall profile. In the larger environment (b), the reflected components lie farther outside the domain and have minimal influence, resulting in a distribution that closely matches the unbounded case (no-reflection condition). **c, d** Successor representations SR_unbounded_(*x*) and SR_bounded_(*x*), computed by summing discounted distributions over *t* = 1 … 30 with *γ* = 0.99. Double-headed arrows indicate the 50% peak level, which demarcates SR place fields. In the short environment (c), broader distributions accumulate to yield a substantially wider SR place field, whereas in the long environment (d), the SR remains tightly localized because boundary reflections contribute only weakly.

**Figure S2.**
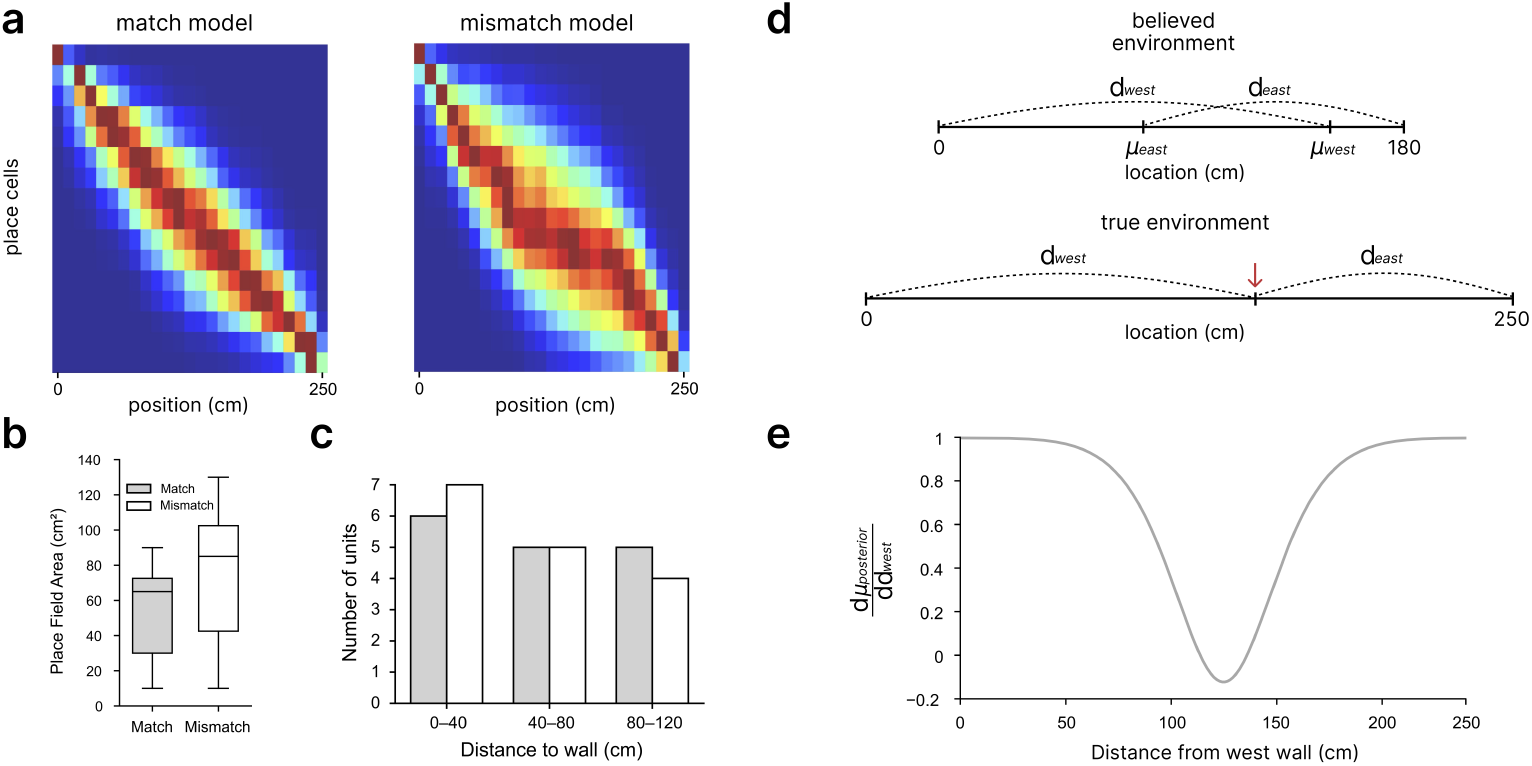
**a**. Heatmaps showing example posterior place fields for 16 units in the match model (left) and the mismatch model (right), with each row representing the spatial tuning of one unit across the environment. **b**. Box plot of place field area for the mismatch and match models. Overall, the mismatch model yields broader place fields than the match model. **c**. Grouped bar plot of place field peak locations for the match (gray) and mismatch (white) models. Peak locations are binned along the length of the environment (0–40 cm, 40–80 cm, 80–120 cm). In the match model, peaks are relatively evenly distributed across bins, whereas in the mismatch model, they are concentrated near the walls. **d**. Schematic showing how opposing visual cues in a mismatched environment influence the expected believed location. **e**. The magnitude of the rate of change of the expected believed location with respect to true location decreases toward the center of the environment, predicting a lower density of place field peaks farther from the wall.

### Why mismatch model’s place fields are broader overall and sparser toward the center

In the main text (Fig. 1d, right), we showed that under geometric mismatch, ideal observer posteriors become anchored relative to positions in the believed environment rather than the true environment, and that this can broaden place fields^55,62^. Fig. S2a illustrates this effect across place cells tuned to evenly spaced believed locations. Each heatmap shows stacked firing rate maps for 16 example units from each of the match and mismatch models. The match model produced narrow fields aligned to the true location, whereas the mismatch model yielded broader fields due to the discrepancy between the believed (180 cm; matching the height of the believed environment in the main text) and true (250 cm; matching the height of environments C and D) environments. This effect was especially pronounced around the center of the environment, where visual cues from both walls are weak and of similar strength, compared with locations nearer either wall, where the visual cue from the closer wall dominates.

Quantification confirmed these differences. Compared to the match model, the mismatch model produced place fields that were broader overall (Fig. S2b) and more concentrated near boundaries (Fig. S2c). These patterns arise because geometric mismatch spreads place fields over a wider area around the center, where visual cues are weaker, while anchoring them closer to walls where visual cues are stronger.

To further examine how place field peak density varies with distance to the wall, we analytically computed the posterior mean location using Bayesian cue combination. This formalizes the intuitive visualization shown in Fig. S2a-c, explaining why place field density decreases with distance.

For simplicity, here we only consider visual input from the west and east walls of a 1D environment (Fig. S2d). The posterior mean (*µ*_post_) combines visual estimates based on the two walls (*µ*_west_ and *µ*_east_), weighted by their relative precisions (*λ*_west_ and *λ*_east_):

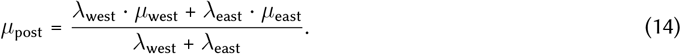

We assume that the visual estimates are well calibrated, such that their means correspond to distances in believed coordinates:

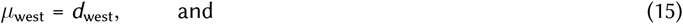

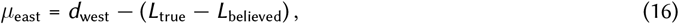

where *d*_west_ is the true distance from the west wall, *L*_true_ is the length of the true environment, and *L*_believed_ is the length of the believed environment. The precision of each cue scales inversely with the fourth power of distance due to the principles of pinhole optics, as explained in Ref. 11:

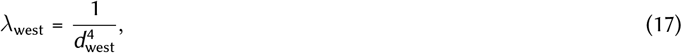

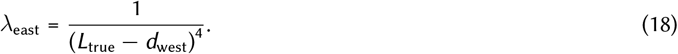

Substituting these into the expression for the posterior mean yields

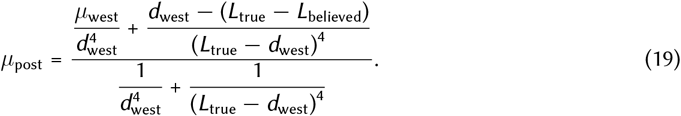

To examine spatial sensitivity, we numerically computed the derivative 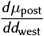. For the numerical evaluation shown in Fig. S2e, we set *L*_believed_ = 180 and *L*_true_ = 250, matching the environment sizes used in Fig. S2a-c, so that *µ*_east_ = *d*_west_ − (*L*_true_ − *L*_believed_) = *d*_west_ − 70.

As shown in Fig. S2e, the magnitude of the derivative decreases toward the center of the environment, and hence farther from the walls, indicating that changes in true location lead to smaller shifts in the posterior mean. Under our assumption that each place cell is tuned to an evenly spaced location in the believed environment, a lower derivative means that more physical distance is required to traverse a unit distance in the believed space. This results in fewer distinct peaks per unit physical distance, that is, a lower density of place fields. Therefore, the observed decrease in the magnitude of the derivative toward the center naturally explains why place field peaks become sparser with distance from boundaries.

In the match model, by contrast, the two wall-based estimates coincide (*µ*_west_ = *µ*_east_ = *d*_west_), so the posterior mean reduces to *µ*_post_ = *d*_west_ and 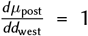 everywhere. Thus, belief changes at a constant rate with physical position, yielding an approximately uniform peak density across distance to the wall, as observed in our simulations.

**Figure S3.**
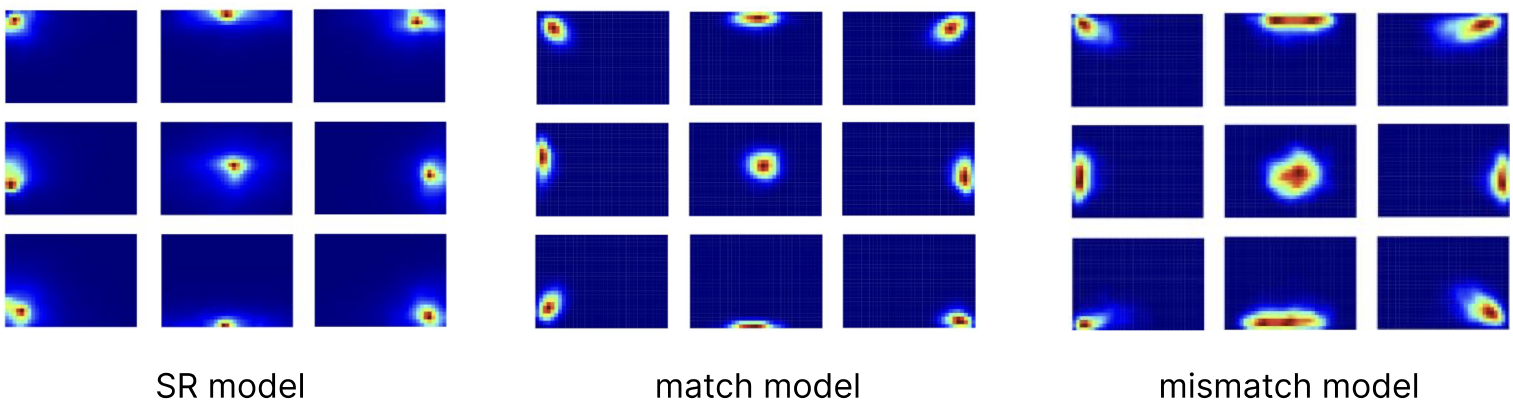
Example units of SR, match, and mismatch models in environment D. Firing rate maps from the SR (left), match (middle), and mismatch (right) models. Each panel shows a single unit, illustrating characteristic differences in field size and shape across models.

**Figure S4.**
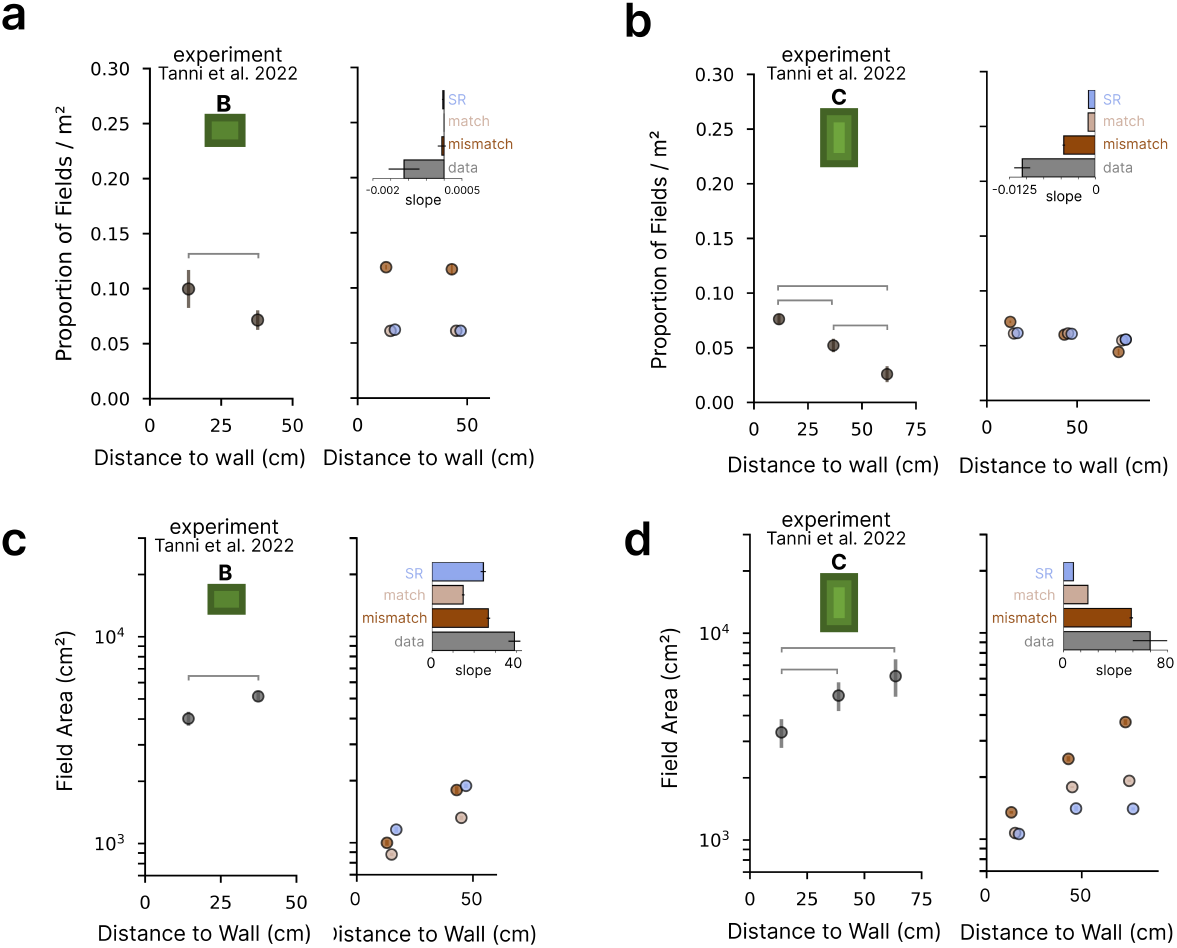
Results for environments B and C. We focused on environment D in the main text because it is the largest and shows the strongest effects of distance to the wall. Here we show environments B and C; environment A is omitted (as in Ref. 24) because it is not large enough to allow comparison across distance bins. In each panel, left shows data and the right shows model prediction. **a**. In environment B, peak density decreased with distance in the data (KW: *H* = 4.81, *p* = 0.028) but not in any model (mismatch: *p* = 0.42; match: *p* = 0.91; SR: *p* = 0.05). **b**. Same as panel **a** for environment C. The mismatch model showed a density decrease similar to the data (data, KW: *H* = 12.5, *p* = 0.0002; mismatch, KW: *H* = 12.5, MW: all *p <* 0.03), whereas the match and SR models showed weaker or inconsistent changes (match, KW: *H* = 9.64, *p* = 0.008, MW: all *p <* 0.04 except 0–30 vs. 30–60 cm, n.s.; SR, KW: *H* = 11.41, *p* = 0.003, MW: all *p <* 0.05), with significant slope differences between models (interaction *p* = 0.023; mismatch vs. match: *p* = 0.042; mismatch vs. SR: *p* = 0.026; match vs. SR: n.s.). **c**. In environment B, average field area increased with distance in the data and all models (data, KW: *H* = 6.82, *p* = 0.009; all models, MW: *p* = 0.008). **d**. Same as panel **c** for environment C. The data showed increasing field area with distance to the wall (KW: *H* = 10.2, *p* = 0.006). The mismatch model recapitulated this increase; the match model showed a weaker rise; and the SR model varied modestly across bins (match and mismatch, KW: *H* = 12.5, *p* = 0.002, MW: all *p <* 0.03; SR, KW: *H* = 9.38, *p* = 0.009, MW: all *p <* 0.03 except 30–60 vs. 60–90 cm, n.s.), with significant slope differences between models (interaction *p* = 0.02; mismatch vs. match: n.s.; mismatch vs. SR: *p* = 0.01; match vs. SR: n.s.)

**Figure S5.**
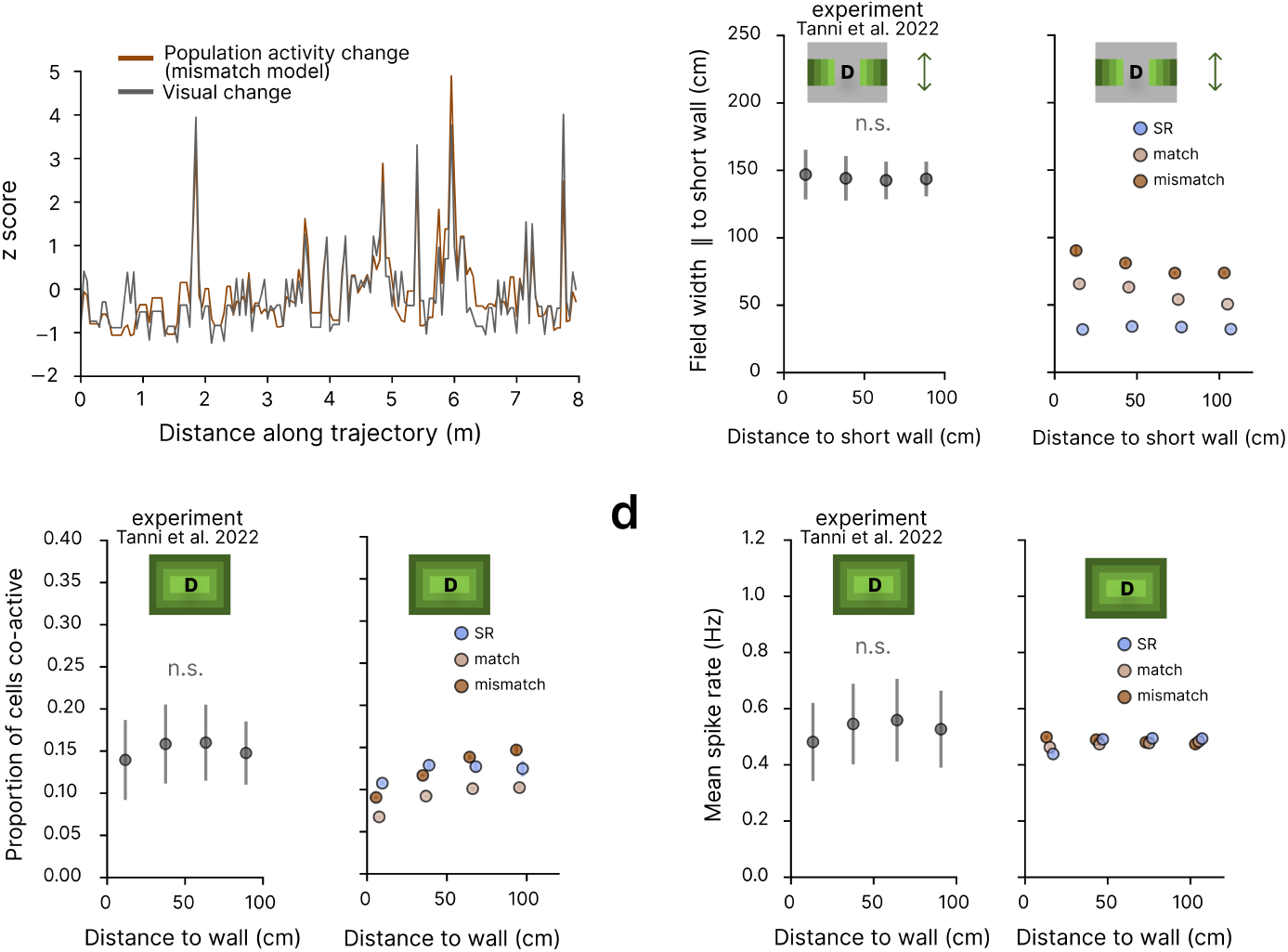
Visual-population coupling and activity modulation in environment D. **a**. Z-scored visual and population activity change along trajectory distance in the mismatch model. **b**. Place field width parallel to the short wall by distance to that wall. Across models, width along this axis was relatively constant with distance, and flatter than along the orthogonal axis (match, KW: *H* = 16.55, *p* = 0.0009, MW: all *p <* 0.05 except 0–30 vs. 30–60 cm, n.s.; mismatch, KW: *H* = 16.07, *p* = 0.001, MW: all n.s. except 30–60 vs. 60–90 cm, *p <* 0.05; SR, KW: *H* = 13.47, *p* = 0.004, MW: all n.s. except 30–60 vs. 90–120 cm and 60–90 vs. 90–120 cm, *p <* 0.05). **c**. Proportion of co-active place cells across spatial bins. Unlike the data, no model showed uniform co-activity. For the match and mismatch models, the proportion increased with distance to wall (match, KW: *H* = 16.14, *p* = 0.001, MW: all *p <* 0.05 except 60–90 vs. 90–120 cm, n.s.; mismatch, KW: *H* = 17.10, *p* = 0.0007, MW: all *p <* 0.05 except 60–90 vs. 90–120 cm, n.s.). In the SR model, the proportion was lower in the outermost bin but stable elsewhere (KW: *H* = 11.45, *p* = 0.01, MW: all n.s. except 0–30 vs. 30–60, 60–90, and 90–120 cm, *p <* 0.05). **d**. Mean firing rate across distance bins. Data showed relatively uniform rates, arising from balanced field density and area. The match and mismatch models showed relatively constant rates (KW: match, *H* = 4.62, *p* = 0.20; mismatch, *H* = 6.23, *p* = 0.10), but for different reasons: elevated rates near boundaries were offset by distance-dependent biases in density and area rather than the density–area balance. In the SR model, rates were lowest in the outermost regions due to smaller field areas and widths, but stable elsewhere (KW: *H* = 10.75, *p* = 0.01, MW: 0–30 vs. 30–60, 60–90, and 90–120 cm, *p <* 0.05; others n.s.).

### Place field properties with lower threshold

Here, place fields are defined as contiguous regions exceeding 30% (instead of the 50% threshold used in the main text) of the peak firing rate. At this lower threshold, SR place fields also exhibit a modest increase in field area and width with distance to the wall (Fig. S6b, Columns 2–4), resembling patterns observed in the ideal observer models and in the data. However, key differences remain: SR field density stays approximately constant across distance and environment sizes, unlike the prominent decrease and increase, respectively, observed in the data and the mismatch model (Fig. S6a). In addition, overall field area still decreases with environment size, unlike the increase observed in the data and the mismatch model (Fig. S6b, Column 1).

**Figure S6.**
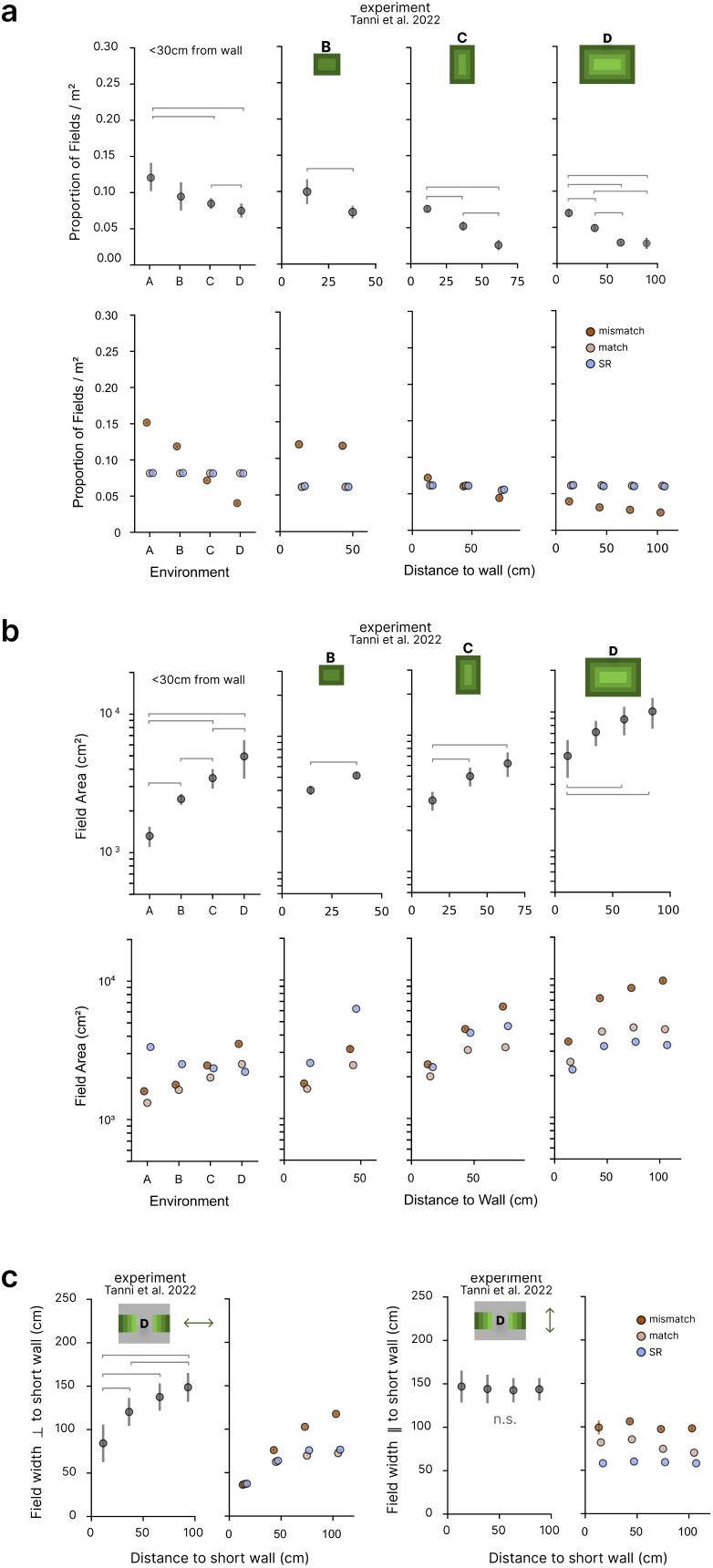
Place field properties when fields are defined as contiguous regions exceeding 30% of the peak firing rate. **a**. Place field peak density across environments and as a function of distance to the wall (top: data, bottom: model results). The results are nearly identical to the main-text analysis at the 50% threshold (Figs. 2 and S4). **b**. Average place field area across environments and as a function of distance to the wall (top: data, bottom: model results). In the SR model, area decreases with increasing environment size. In environment D, a prominent increase with distance is observed only in the mismatch model. **c**. Field width orthogonal to the short wall (left) and parallel to the short wall (right) as a function of distance to the short wall. The orthogonal width increases most strongly in the mismatch model, while the parallel width remains roughly constant across models.

### One-step SR model

We have also considered a one-step variant of the SR (“one-step SR model”). It is defined simply by truncating the series in Eq. 5 at *τ* = 1, i.e.,

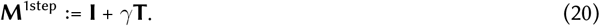

Here we show that the one-step SR model yields the same place fields as those based on the predictive distribution of the Bayesian ideal observer model with the transition matrix

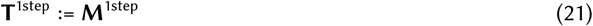

in the limiting case that ignores perceptual uncertainty (i.e., assumes noiseless, uniquely identifying observations). Formally, it treats

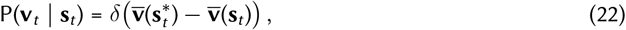

where 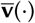 is the expected visual input at a given state, 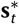 is the true state, and the visual input is assumed to match the expectation (since we are ignoring perceptual uncertainty), i.e., 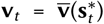. Substituting this into the measurement step (Eq. 2) yields:

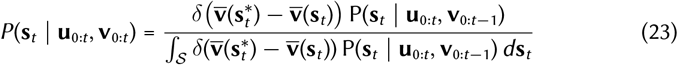

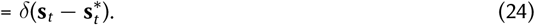

In other words, if we ignore perceptual uncertainty, as soon as sensory measurement is made, the posterior collapses onto the true state, which serves as the dynamic prior for the next time step. Substituting this uncertainty-free dynamic prior into the transition step (Eq. 1) shows that the resulting predictive distribution reduces to the transition probability:

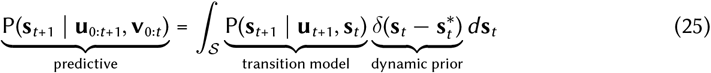

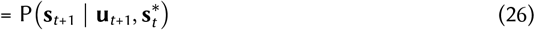

The average of this transition probability across time steps, as taken when computing the SR place fields, is proportional to the transition matrix (see Eq. 8):

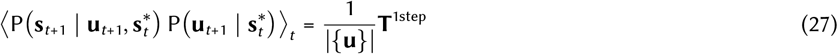

Therefore, the one-step SR model is equivalent to the Bayesian ideal observer without perceptual uncertainty, with **T**^1step^ := **I**+*γ***T**, and thus yields identical place fields. It produced results qualitatively similar to the multi-step SR model, including its failure to reproduce key trends observed in the data (see also Fig. 2c–h).

**Figure S7.**
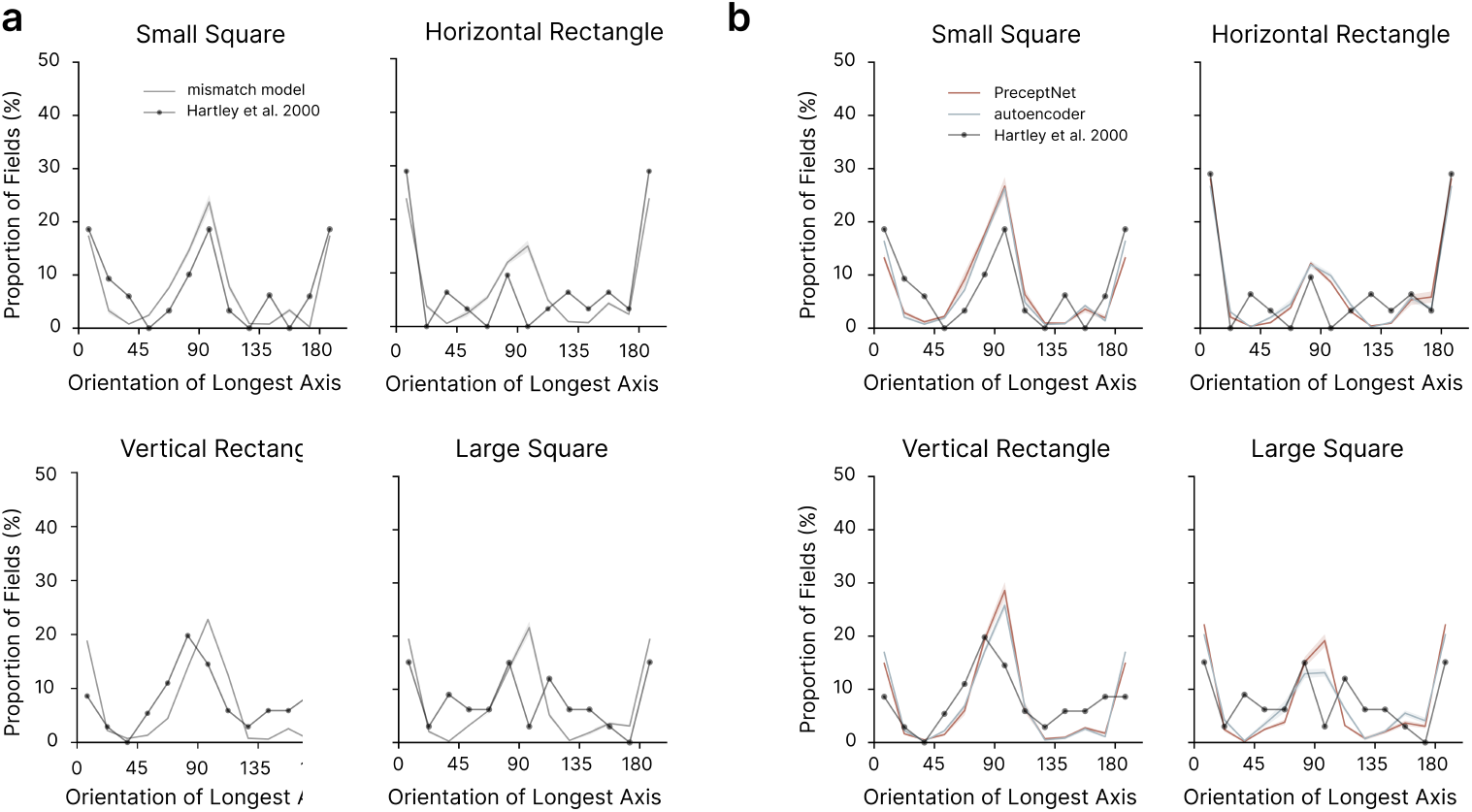
Place field orientation across environments. **a**. Proportion of fields by the orientation of their longest axis for the mismatch model, compared to the experimental data^19,35^. **b**. Corresponding distributions for PreceptNet and the autoencoder models across four environments (small square, large square, vertical rectangle, and horizontal rectangle).

### Cue-conflict after the agent’s 180° rotation in a geometrically mismatched environment

In the main figure (Fig. 5d), we showed cue-conflict steps involving 90° rotations. Here, we extend this analysis to include cue conflict arising from 180° rotations. For this, we specifically selected steps where the agent was in either the top or bottom third of the environment (Fig. S8b). This choice emphasizes cases where cue conflict is especially pronounced. As in the 90° case, no-conflict steps were defined as time steps spent in these regions without rotation.

The KL divergence from the ideal observer was smaller for the posterior decoded from PreceptNet (“PreceptNet’s posterior”) than with the autoencoder’s posterior in the cue-conflict steps (Fig. S8c; PreceptNet: 8.19 *±* 0.58; autoencoder: 22.66 *±* 0.61; paired t-test, *t*(4) = −13.83, *p* = 0.0002). This pattern also held in no-conflict steps (Fig. S8c; PreceptNet: 3.19 *±* 0.11; autoencoder: 4.29 ± 0.03; paired t-test, *t*(4) = −13.41, *p* = 0.0002), though the difference between models was smaller (cue-conflict steps: 14.46 ± 1.05, no-conflict steps: 1.10 ± 0.08; paired t-test, *t*(4) = 13.07, *p* = 0.0002). Notably, 180° rotations induce a substantial mismatch between the dynamic prior and the visual likelihood. Thus, the autoencoder’s posterior jumps abruptly as it relies heavily on the visual likelihood, while PreceptNet’s posterior shifts more smoothly, indicating greater use of the dynamic prior (Fig. S8a).

**Figure S8.**
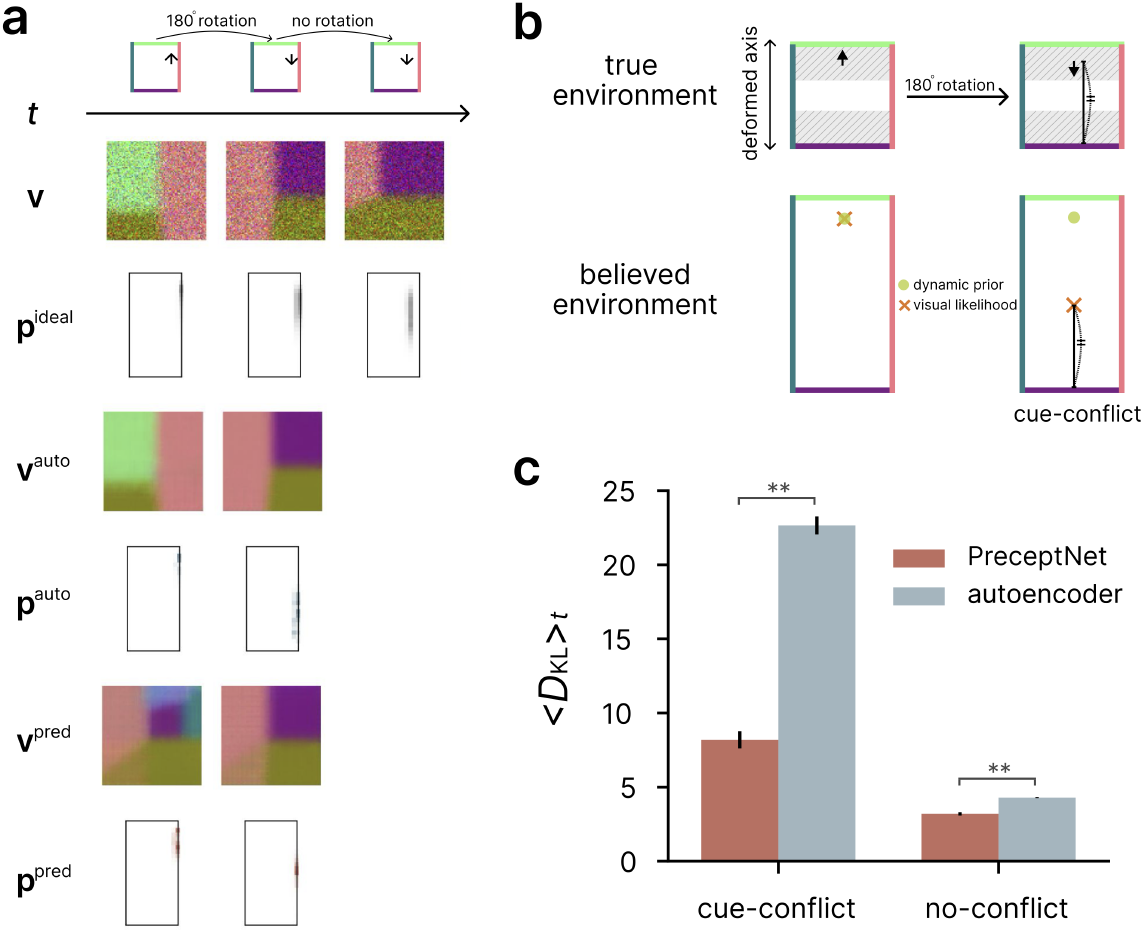
Cue-conflict after the agent’s 180° rotation in a geometrically mismatched environment. **a**. Example time steps in the small square environment with cue-conflict. **b**. Schematic illustration of 180° rotation cue-conflict trials. The agent rotates 180° in the true environment (top), causing a mismatch between its dynamic prior and visual likelihood in the believed environment (bottom). **c**. Average *D*_KL_ between decoded and ideal posteriors in cue-conflict and no-conflict steps.

### Why prediction encourages posterior representations

We show that each layer of the RNN can be made to correspond to successive steps of Bayesian filtering, as depicted in Fig. 6b. We further suggest that the training objective determines which latent variables are useful to represent: the autoencoding objective favors representation of the visual likelihood, whereas the predictive objective encourages additional representation of the predictive distribution and the posterior.

#### Setup and notation

As introduced in Bayesian ideal observer model, the ideal observer separates localization into three computational stages: extracting what the current image alone implies (visual likelihood), combining it with the previous belief (measurement step), and propagating the resulting belief through self-motion (prediction step).

To express these relations in matrix notation, we discretize the state space^63^. Let **s**_*t*_ ∈ *{*1, …, *S}* denote the latent state (pose in our case) and 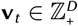 the observation (visual inputs with *D* pixels in our case) at time *t*.

To compute the visual likelihood, we assume Poisson-distributed visual input, which approximates input from a retinotopically arranged layer of neurons. Then the environment’s observation model can be expressed as a fixed matrix 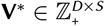 whose **s**-th column 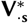 is the expected image at state **s**. The generative distribution is

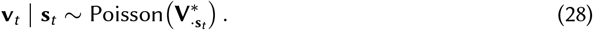

Applying Bayes’ rule to this generative model yields the vision-only posterior (normalized visual likelihood), denoted 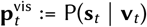). The next two stages compute the posterior, 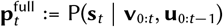, and the predictive distribution, 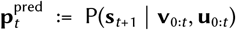 := P(**s**_*t*+1_ **v**_0:*t*_, **u**_0:*t*_) (Eqs. 1 and 2). This ideal observer can be written compactly as:

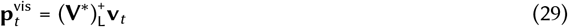

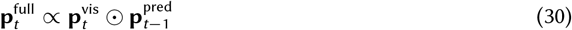

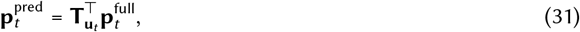

where 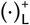 denotes the left pseudoinverse, ⊙ the element-wise (Hadamard) product, and **T**_**u**_ the transition matrix conditioned on control signal **u**. Likewise, the network’s operation can be written as:

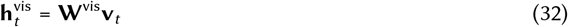

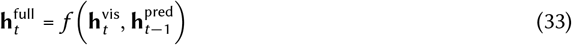

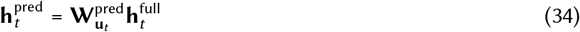

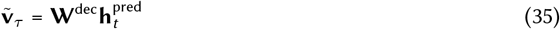

where **h**^vis^, **h**^full^, and **h**^pred^ are termed visual, posterior, and predictive hidden states, paralleling the names of the corresponding stages of Bayesian filtering, and 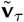 is the visual input predicted for the target time step *τ* . We ignore nonlinearities in Eqs. 32 and 34 as they are unnecessary if we make a simplifying assumption that the network has enough capacity, as shown below. In our study, the autoencoder and PreceptNet have identical structures, which is mirrored in this simplified analysis. The only difference is the target time step: the autoencoder’s target is the current visual input (*τ* = *t*), whereas PreceptNet’s is the next visual input (*τ* = *t* + 1). Finally, both the autoencoder and PreceptNet are trained to minimize the Poisson NLL loss:

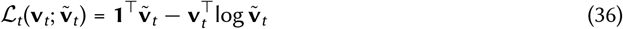

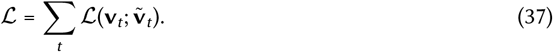

#### Autoencoder: reconstruction objective leads to visual likelihood encoding

As described above, the autoencoder is trained to reproduce the current input. The true conditional mean of the current input is:

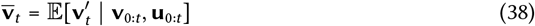

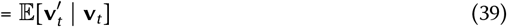

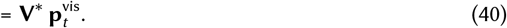

Then, in the limit of a large data set, minimization of the loss (Eq. 37) requires the decoded output 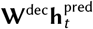 at each time step *t* to satisfy

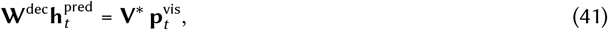

as obtained by optimizing Eqs. 36 and 37 after substituting Eqs. 29 and 35. One such solution for the weight matrices in Eqs. 32-35 is to set

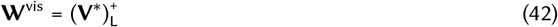

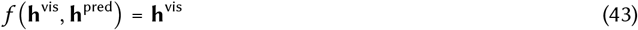

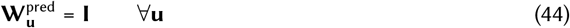

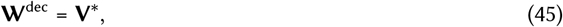

which yields, for each *t*, all hidden states matching the vision-only posterior:

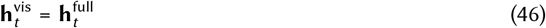

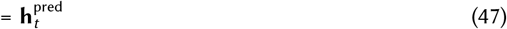

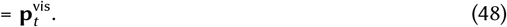

These results reveal a crucial insight: the autoencoding objective (Eq. 37) can be satisfied by simply inverting the generative process of the visual input in the encoding step (as in Eq. 42 for Eq. 32), the first step in the chain of operations (Eqs. 32-35). This renders the rest of the operations redundant: it is enough for the subsequent operations to simply preserve the result of the first operation. Notably, both history dependence (conveyed by 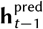 in Eq. 33) and control dependence (conveyed by **u**_*t*_ in Eq. 34) can be ignored (Eqs. 43 and 44). This makes the loss at each time step independent of variables at other time steps in Eq. 37.

Also as a consequence, none of the hidden layers (**h**^vis^, **h**^full^, or **h**^pred^) consistently represents the posterior, **p**^full^, which is distinct from what they represent, **p**^vis^ (Eq. 30). This is consistent with our numerical findings: **p**^vis^ can be decoded from **h**^pred^ better than the posterior **p**^full^ or the predictive **p**^pred^ (Fig. 6c, top).

#### PreceptNet: predictive objective leads to posterior encoding

PreceptNet is trained to predict the next input. The true conditional mean of the next input is:

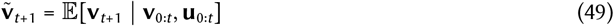

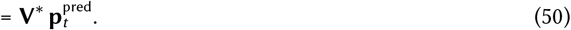

Note that the last term involves **p**^pred^ rather than **p**^vis^, unlike Eq. 41. Then, minimization of Eq. 37 requires, paralleling Eq. 41,

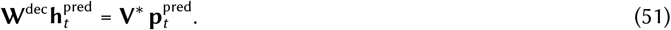

One such solution for Eqs. 32-34 is to set

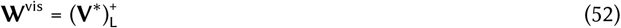

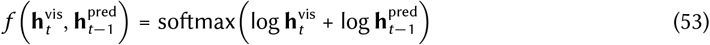

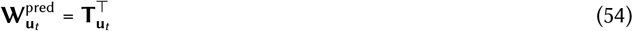

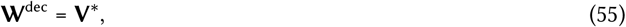

which yields

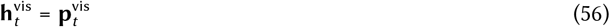

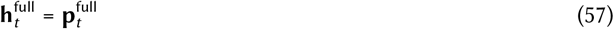

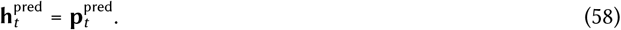

Note that, unlike the autoencoding objective, the predictive objective cannot be satisfied by applying a fixed operation to the visual input alone, except in degenerate cases where the next visual input is fully determined by the current visual input, such as when the same visual input distribution is expected at every time step regardless of control. Rather, it requires incorporation of information about the control (**u**_*t*_ in the right hand side of Eq. 54) and history information (which shapes the ideal observer’s predictive distribution; note that 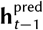 is in the right hand side of Eq. 53 but not Eq. 43). While the sequential correspondence between the stages of Bayesian filtering and the hidden states of PreceptNet in Eqs. 56-58 is a sufficient but not necessary condition for Eq. 51, our numerical results nevertheless show just such a correspondence (Fig. 6c, bottom).

Notably, this couples the loss of PreceptNet across time steps through **h**^pred^ (in Eq. 37), unlike the autoencoder. This leads to different representations being learned at each layer, as illustrated by the case of ambiguous states, discussed next.

#### Ambiguous states

In the main text, we use ambiguous states, the states that share the same expected visual input, to illustrate the difference between the autoencoder and PreceptNet. Here we provide an analytical argument for the difference.

For the autoencoder, when the agent is in one of ambiguous states, the decoded posterior is characteristically broad. This can be understood from the fact that in Eq. 29, ambiguous states make **V**^∗^ contain identical columns (corresponding to identical visual inputs across different ambiguous states) and hence become rank-deficient. In this case, we can separately collect a set of identical columns, 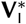, and the rest, 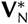, as submatrices and let

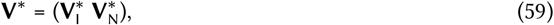

and make the corresponding separation as

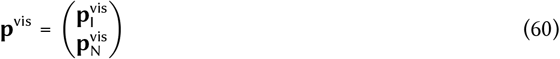

Here, the pseudoinverse chooses the minimum-norm solution, making elements of 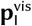 identical. This coincides with the ideal observer’s vision-only posterior in our setting: states with the same expected visual inputs must have the same visual likelihood. This, in turn, makes the decoded posterior broader (Fig. 4b). In other words, the autoencoder cannot distinguish between ambiguous states.

In contrast, PreceptNet overcomes two challenges involving ambiguous states. First, as with the autoencoder, the state **s**_*t*_ at a particular time step *t* may be ambiguous, i.e., may share identical visual input **v** with another state visited at another time step, **s**_*t*_*′* . This will yield the same hidden state 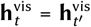. However, as long as their past trajectories differ, the previous predictive hidden states will be different, i.e., 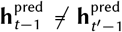, which results in different posterior hidden states, 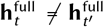, and in turn, different predictive hidden states, 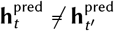. Therefore, representations of distinct posteriors can be learned. In other words, PreceptNet can distinguish between ambiguous states based on the history (see also Fig. 6a, bottom).

Second, the next sensory input may be the same (**v**_*t*+1_ = **v**_*t*_*′*_+1_), even when both the past trajectories and the next states are distinct (**s**_0:*t*_ **s**_0:*t*′_ and **s**_*t*+1_*≠* **s**_*t*′+1_). In this case, generally, the two predictive states are different 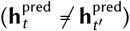, yet they will need to predict the same sensory input. This is possible due to the very ambiguity of the states. That is, if we make separations similar to Eq. 60:

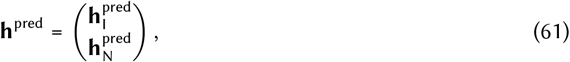

the same visual stimulus can be predicted from multiple different values of 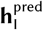 as long as the sum of elements within the ambiguous states, 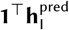, remains the same.

Since the predictions at different time steps are connected by **h**^pred^ in PreceptNet (unlike in the autoencoder), the elements of 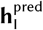 at one time step eventually contribute to the loss computed with 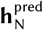 at another time step, as long as the unambiguous states are reachable from the ambiguous states, and vice versa, as in the experiments considered in this study. This allows the weights of PreceptNet to be optimized to consider history in predicting the next sensory input.

**Figure S9.**
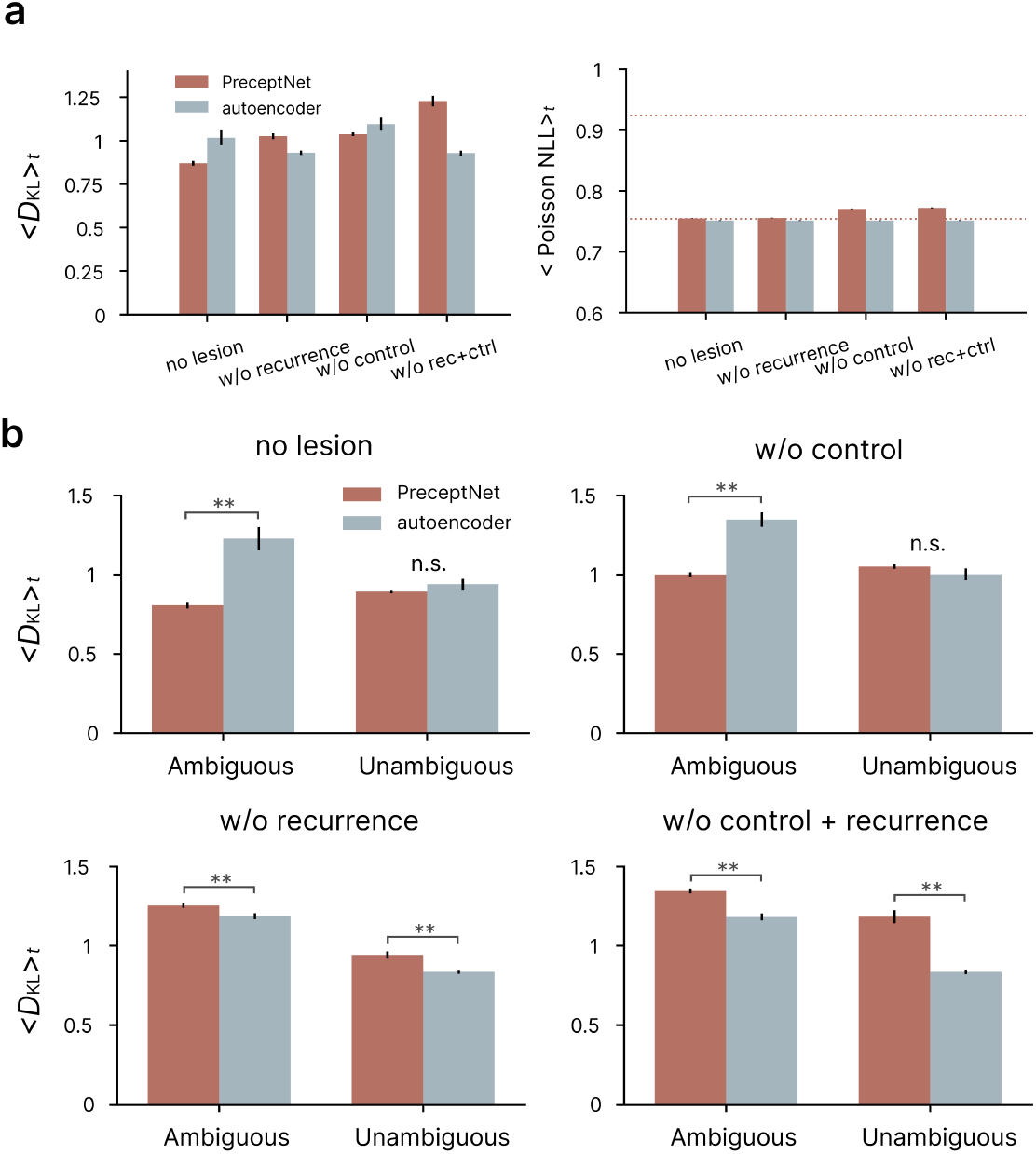
Lesion effects on posterior encoding and training objectives. **a**. *Left:* Average *D*_KL_ between the decoded posterior and the ideal observer posterior in intact networks and networks with recurrence, control, or both inputs ablated. Maximum *D*_KL_ values estimated from shuffled data were 3.45 (PreceptNet) and 1.64 (autoencoder). PreceptNet without lesion achieved the minimum *D*_KL_, reflecting contributions of recurrence and control. *Right:* Average loss under each model’s training objective: Poisson negative log-likelihood (NLL) between predicted and actual visual input for PreceptNet, and reconstruction loss for the autoencoder. For PreceptNet, losses were: no lesion, 0.755 *±* 0.001; w/o recurrence, 0.755 *±* 0.001; w/o control, 0.770 ± 0.001; w/o both, 0.772 ± 0.001. For the autoencoder, all conditions yielded 0.752 ± 0.001. Dotted lines indicate the minimum achievable loss (0.754) and the maximum loss measured with shuffled data (0.819) for PreceptNet. See also Fig. 6d for the differences between the lesioned and intact networks. **b**. Average *D*_KL_ between the decoded and ideal observer posteriors, shown separately for ambiguous and unambiguous states in intact networks and after ablating recurrence, control, or both inputs, as in panel **a**. For PreceptNet, recurrence was especially critical under ambiguous inputs: removing recurrence abolished its advantage over the autoencoder. By contrast, control lesions moderately increased *D*_KL_ across both ambiguous and unambiguous states. The combined lesion produced the largest overall impairment. The autoencoder showed no significant changes across conditions.

**Figure S10.**
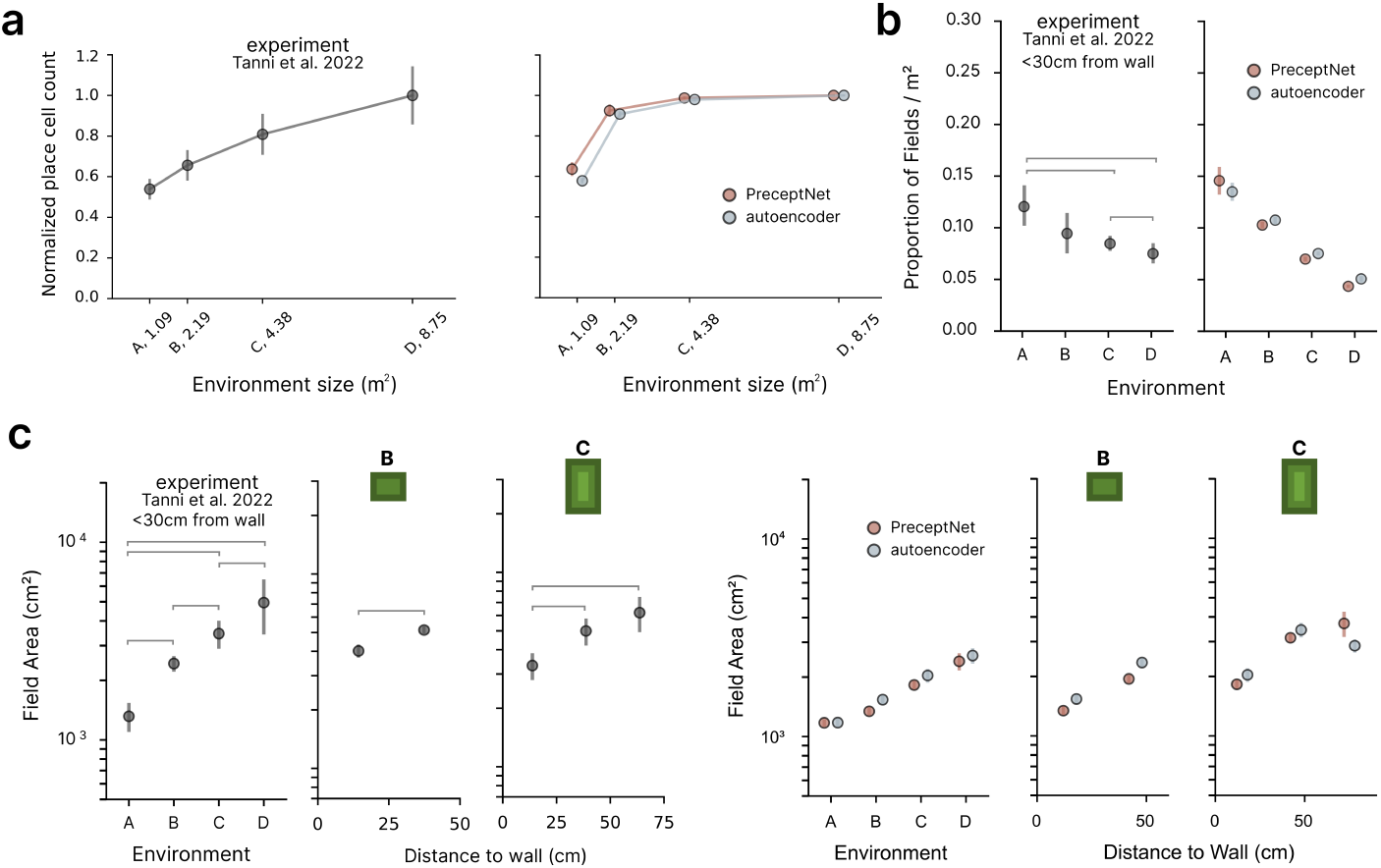
Place cell properties in PreceptNet and autoencoder models across environment sizes and spatial positions. **a**. Number of active place cells (firing rate *>* 1 Hz) increased with environment size in both PreceptNet and autoencoder. **b**. Place field density decreased with environment size in both PreceptNet and autoencoder (both, KW: *H* = 17.86, *p* = 0.0005, MW: all *p <* 0.05). **c**. Average place field area increased with environment size for both models (PreceptNet, KW: *H* = 17.86, *p* = 5 *×* 10^−4^; autoencoder, KW: *H* = 17.58, *p* = 5 *×* 10^−4^, MW: all pairwise *p <* 0.05). Within environments B and C, PreceptNet showed an increasing trend with distance to the wall (B, MW: *p* = 0.01; C, KW: *H* = 10.5, *p* = 0.005, MW: all *p <* 0.03, except 30–60 vs. 60–90, n.s.), whereas the autoencoder showed a significant drop in the farthest bin (B, MW: *p* = 0.008; C, KW: *H* = 11.0, *p* = 0.003, MW: all *p <* 0.04). See also Fig. 5g–i.

